# Spatiotemporal Genomic Profiling of Intestinal Metaplasia Reveals Clonal Dynamics of Gastric Cancer Progression

**DOI:** 10.1101/2023.04.10.536195

**Authors:** Kie Kyon Huang, Haoran Ma, Tomoyuki Uchihara, Taotao Sheng, Roxanne Hui Heng Chong, Feng Zhu, Supriya Srivastava, Su Ting Tay, Raghav Sundar, Angie Lay Keng Tan, Xuewen Ong, Minghui Lee, Shamaine Wei Ting Ho, Tom Lesluyes, Peter Van Loo, Joy Shijia Chua, Kalpana Ramnarayanan, Tiing Leong Ang, Christopher Khor, Jonathan Wei Jie Lee, Stephen Kin Kwok Tsao, Ming Teh, Hyunsoo Chung, Jimmy Bok Yan So, Khay Guan Yeoh, Patrick Tan, Singapore Gastric Cancer Consortium

## Abstract

Intestinal metaplasia (IM) is a pre-malignant condition of the gastric mucosa associated with increased gastric cancer (GC) risk. We analyzed 1256 gastric samples (1152 IMs) from 692 subjects through a prospective 10-year study. We identified 26 IM driver genes in diverse pathways including chromatin regulation (*ARID1A*) and intestinal homeostasis (*SOX9*), largely occurring as small clonal events. Analysis of clonal dynamics between and within subjects, and also longitudinally across time, revealed that IM clones are likely transient but increase in size upon progression to dysplasia, with eventual transmission of somatic events to paired GCs. Single-cell and spatial profiling highlighted changes in tissue ecology and lineage heterogeneity in IM, including an intestinal stem-cell dominant cellular compartment linked to early malignancy. Expanded transcriptome profiling revealed expression-based molecular subtypes of IM, including a body-resident “pseudoantralized” subtype associated with incomplete histology, antral/intestinal cell types, *ARID1A* mutations, inflammation, and microbial communities normally associated with the healthy oral tract. We demonstrate that combined clinical-genomic models outperform clinical-only models in predicting IMs likely to progress. Our results raise opportunities for GC precision prevention and interception by highlighting strategies for accurately identifying IM patients at high GC risk and a role for microbial dysbiosis in IM progression.

## Introduction

Gastric cancer (GC) is a major cause of global cancer burden [1]. Despite an overall decline in age-adjusted incidence, GC still ranks fifth in incidence and fourth in mortality [2]. GC generally carries a poor prognosis as diagnosis is often made at late disease stages, and in younger patients (<50 years) increasing GC incidence in the stomach body and cardia has been reported [3, 4]. In countries with high GC prevalence such as Japan and South Korea, population screening has resulted in improved outcomes due to early detection [5]. However, in many countries such as Singapore where GC incidence is moderate, population screening is not cost-effective [6]. There is thus a need to better understand the pathogenesis of GC to guide precision prevention efforts.

The stomach is a complex organ with distinct anatomical regions (antrum, body, and cardia) harbouring different cell types and functionalities [7]. GC can arise in any of these regions through the interaction of genetic and environmental factors including *Helicobacter pylori* (Hp) infection [8]. An important step in GC carcinogenesis is intestinal metaplasia (IM) [9], a pre-malignant condition where cells lining the stomach are replaced by cells with characteristics similar to the small intestine. However, although IM patients have increased GC risk (6 fold; [10]) it remains unclear if IM cells represent direct precursors of malignancy, or if the presence of IM reflects bystander tissue damage caused by Hp and chronic inflammation [11, 12, 13]. Some groups have proposed that IM cells, being post-mitotic and differentiated, are unlikely to cause cancer [11, 13], and that GC may emerge from other gastric epithelial stem cell populations due to the ability of stem cells to self-renew and survive for prolonged periods [11, 13]. Evidence also suggests that IM are heterogenous between and within patients. For example, IMs can exhibit either “complete” histology (Type I) with small intestinal-type mucosa and mature absorptive cells, goblet cells and brush borders, or “incomplete” histology (Type III) with colonic epithelium and columnar ‘intermediate’ cells in various stages of differentiation, with the latter associated with higher GC risk [14]. In many patients, IM initially occurs in the gastric antrum expanding to the body and cardia, and GC risk is higher when IM involves both the antrum and body/cardia compared to IMs involving the antrum only [15]. Besides IM, other variants of metaplasia have also been reported in the gastric body, such as “pyloric metaplasia” or Spasmolytic Polypeptide-Expressing Metaplasia (SPEM), a metaplastic mucous cell lineage with phenotypic characteristics of deep antral gland cells [12]. To date, only a handful of studies have examined genomic and molecular features of IM [16, 17, 18].

A comprehensive molecular study of IM is thus needed to better understand IM molecular landscapes, inter- and intra-patient heterogeneity, and relationships between IM and GC at the genomic and clinical level [16]. Here, we performed a comprehensive analysis of IMs sampled from a prospective clinical study, leveraging high-depth targeted DNA sequencing, transcriptome sequencing, and recently developed single-cell and spatial transcriptomic platforms. We identified novel IM driver genes, differentially expressed subtypes, and changes in cellular compositions linked to the expansion of specific stem cell communities. We also discovered a potential role for microbial dysbiosis in the pathogenesis of a subset of IMs.

## Results

### Study Design and Datasets

The Gastric Cancer Epidemiology Program (GCEP) is a prospective multi- center longitudinal cohort study, monitoring 2980 Chinese participants aged ≥50 from 2004 to 2015 [19]. GCEP subjects underwent screening gastroscopies with standardised gastric mucosal sampling at multiple stomach regions (antrum, body, cardia) and surveillance endoscopies at years 3 and 5. At study conclusion, 82% of subjects had completed 5 years of follow-up, collectively representing 11157 person-years of surveillance (Supplementary Figure S1).

We performed high-depth (>1000x) targeted DNA sequencing of 277 cancer genes on 1217 endoscopic biopsies from 682 unique subjects (1119 samples from 644 subjects with IM; 98 samples from 38 control subjects without IM) (Supplementary Table 1). To enable intra-patient (ie within-patient) comparisons, we profiled samples from multiple stomach sites (antrum: n=642; body: n=274; cardia: n=265). A subset of samples were matched from the same subjects across time, enabling longitudinal comparisons from subjects who i) developed dysplasia during their course of observation (n=64), ii) had concurrent dysplasia (n=93) or iii) exhibited dysplasia regression (n=98)) (**Figure 1A**). Selected IM samples with appreciable median variant allele frequencies (VAFs) were analysed by whole-genome sequencing (WGS, n=5) to assess mutational counts and signatures. At the transcriptomic level, we performed bulk RNA-sequencing on 183 GCEP samples, including normal (n=46) and IMs (n=137) from multiple sites (antrum: n=55; body: n=66; cardia: n=62).

**Figure 1.**
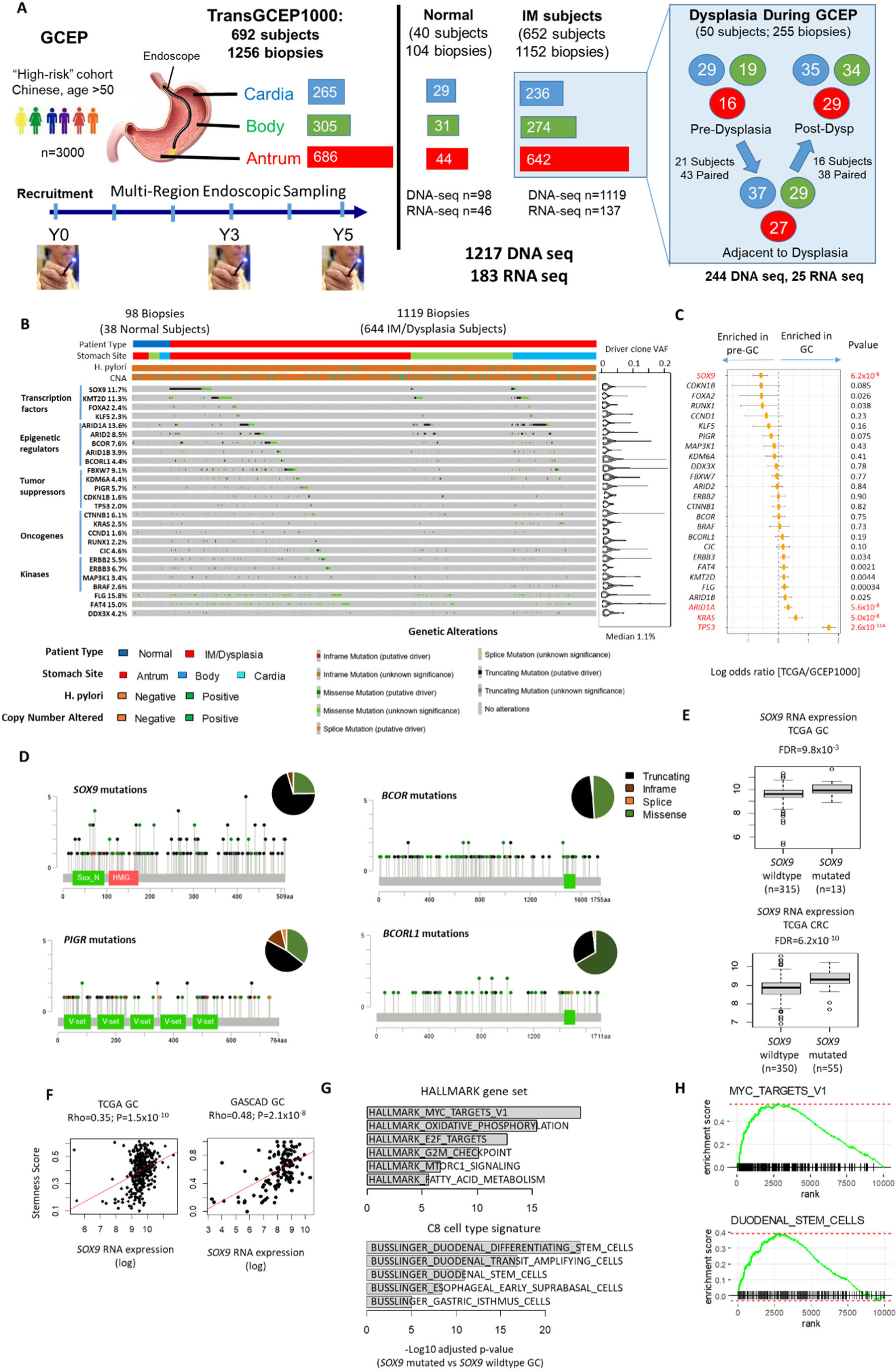
Genomic profiles of gastric pre-malignancy. (A) Overview of the TransGCEP1000 translational study. 1256 gastric biopsies from multiple stomach sites were analyzed from 692 GCEP subjects. (Right) A subset of samples were longitudinally matched from the same subject, from either pre-dysplasia to dysplasia (adjacent) or dysplasia (adjacent) to post-dysplasia where regression was observed. (B) Oncoplot showing predicted IM driver genes. (Right) Violin plots indicate median VAFs of detected somatic mutations. (C) Log odds ratios of driver gene mutation frequencies in TCGA (GC) vs TransGCEP1000 (pre-malignancy). Left shifted genes are mutated more frequently in pre-malignancy, while right-shifted genes are mutated more frequently in GC. (D) Lollipop plot showing distributions and categories of protein altering mutations in *SOX9*, *PIGR*, *BCOR* and *BCORL1*. Pie charts indicate the percentage of different types of non- synonymous mutations. (E) Boxplot comparing *SOX9* RNA expression levels in *SOX9*-mutated and *SOX9*-wildtype GCs (upper) and colorectal cancers (lower). (F) Correlation between *SOX9* expression with TCGA mRNA stemness score in TCGA GCs (left) and a separate cohort (GASCAD, right) of GC samples. (G) Geneset enrichment analysis of *SOX9* mutated vs *SOX9* wildtype GCs using the Hallmark database (upper) and Busslinger et al dataset [35] (lower). (H) GSEA plots showing enrichment of MYC target V1 pathway genes and duodenal stem cell signatures in *SOX9* mutated GCs.

To complement the GCEP data, we further generated a) whole-exome sequencing (WES) data of 28 cases of concurrent normal, dysplasia and early GC from South Korea, b) single-cell RNA-sequencing (scRNA-seq) from 10 patients with antral IM and 4 patients with body/cardia IM to survey tissue ecologies, and c) Nanostring DSP spatial profiles of 8 patients whose antral sections contained histologically normal, IM, GC, lymphoid aggregates, and stromal regions, representing 480 regions of interest (ROIs) and 76 CD45-segmented areas of illumination (AOIs).

### Driver gene landscape of gastric pre-malignancy

We sequenced each GCEP sample across 277 human genes associated with gastrointestinal (GI) cancer and other GI conditions (Supplementary Table 2). Average coverage was 1046x to confidently identify small clonal events. We identified 23,575 somatic mutations across the 1217 samples with a median VAF of 1.0% (range 0.075% to 36.7%; compared to paired blood samples). Consistent with previous reports [16], the IM mutation rate was significantly higher compared to normal gastric samples within the genomic regions analysed (1.97 Mb; (9.6 vs 1.8 mutations/Mb; Wilcoxon test p<2.2x10^-16^) (Supplementary Figure 2A). Mutation rates correlated with subject age (r=0.26, Pearson’s correlation test, p-value<2.2x10^-16^) with the highest mutation rates in the antrum (12.7, 2.5, 7.6 mutations/Mb in antrum, body and cardia, Kruskal-Wallis test p<2.2x10^-16^) (Supplementary Figure 2A, B). Most IM samples exhibited mutational signatures associated with SBS1 (aging; 97% of IM biopsies), with smaller contributions of SBS18 (oxidative stress; 3.2%), SBS5 (clock-like signature; 2.6%), SBS3+8 (homologous recombination; 1.1%) and SBS17 (unknown etiology; 1.1%) (Supplementary Figure 2C, D). Expanded WGS analysis of 5 IMs with elevated VAFs (average genome coverage 69.4X) confirmed the presence of SBS1, SBS18, and SBS17 signatures (Supplementary Figure 2E, F), with higher overall contributions of SBS18 and SBS17 likely due to the larger number of somatic SNVs (median 7327; range 3346 to 16829) recovered from WGS.

Using dNdScv [20] to identify genes under positive selection, we identified 26 candidate driver genes (q<0.15) (**Figure 1B**). These included credentialed oncogenes (e.g. *KRAS*, *ERBB2*, *ERBB3, BRAF*) and tumor suppressors (e.g. *ARID1A*, *TP53*) including *FBXW7* which we previously reported [16]. Most oncogene mutations in *KRAS*, *ERBB2*, *ERBB3* and *BRAF* were missense mutations (213/224; 95.1%), including activating mutations at *KRAS* G12 (n=7) and G13 (n=7), *ERBB2* S310 (n=15) and R678 (n=9) and *ERBB3* V104 (n=11) (Supplementary Figure 2G). *BRAF* mutations occurred in regions other than position V600 (G466; n=2, G469; n=3 and D594; n=4). *ARID1A* mutations occurred as missense (n=57), nonsense (n=53), splice-site (n=9) and indel (n=97) mutations in 13.6% of samples (165/1217). However, in contrast to GC where most *ARID1A* indel mutations occurred at microsatellites in microsatellite instability-high (MSI-H) tumors, *ARID1A* indels in the GCEP samples localized largely to microsatellite-free regions (GCEP: 85/97; 87.6% vs TCGA 25/80; 31.2%, Fisher-test, odds ratio 15.3, p-value 7.2x10^-15^). Besides *ARID1A*, we also observed mutations in the related SWI/SNF subunits *ARID1B* and *ARID2* in 3.9% (48/1217) and 8.5% (103/1217) of samples. Many of these driver events were present at relatively low VAFs (median 1.1%, range 0.093% to 21.0%) consistent with pre-malignant gastric samples harbouring multiple small and genetically diverse clones. Notably, *TP53* was mutated in only 2.0% (24/1217) of premalignant samples compared to 48.9% of GCs (TCGA 213/436; Fisher-test, odds ratio 0.02, p-value<2.2x10^-16^) (**Figure 1C**), suggesting that *TP53* mutations are likely to occur later in gastric tumorigenesis after IM onset.

Other notable driver genes included *SOX9*, *PIGR*, *BCOR*, *BCORL1* and *KLF5* (**Figure 1D**, Supplementary Figure S2G). Mutations in *SOX9* (discussed below) and *PIGR* were mostly truncating mutations. *PIGR* encodes a polymeric immunoglobulin receptor and *PIGR* mutations have be reported previously in inflammatory bowel disease and GC ([21, 22, 23]). *BCOR* (BCL6 corepressor) and *BCORL1* (a BCOR homolog; BCL6 corepressor-like 1) encode transcriptional corepressors forming part of the PRC1.1 variant polycomb repressive complex 1 [24]. *KLF5* displayed missense mutations around codons 301-307, consistent with previous reports of driver mutation patterns in this gene [25] (Supplementary Figure S2G). Certain genes were mutated multiple times in the same sample (*ARID1A* 35/165; *ARID2* 16/103, *SOX9* 25/142), consistent with either two-hit inaction or convergent evolution in different clones.

Besides coding exons, our sequencing panel also targeted Hp genes and ∼5000 SNP sites distributed throughout the genome, allowing us to infer Hp infection status and copy number alterations (CNAs). High Hp burden (>10X coverage) was observed in 6.1% of biopsies from IM subjects (68/1119) compared to 1.0% of normal samples (1/98; Fisher test p-value 0.037). Hp was more readily detected in body/cardia biopsies of IM patients compared to antrum IM biopsies (38/487; 7.8% vs 30/632; 4.7%, Fisher test p-value 0.043) (see Discussion). (Supplementary Figure 3A). 4 significant CNA regions were identified (7q36 and 8q24 amplifications, 8p23 and 11p15 deletions) (Supplementary Figure 3A and 3B). Notably, samples with SBS18 signatures exhibited higher mutation rates (Wilcoxon test p-value 9.1x10^-3^) and CNAs (Fisher test p-value 2.7x10^-3^ for GATK; 9.1x10^-3^ for ASCAT) consistent with enhanced oxidative stress contributing to genomic instability (Supplementary Figure 3C).

Several GCEP1000 driver genes have been reported to be involved in intestinal stem cell homeostasis (eg *SOX9*, *ARID1A*, *FBXW7*, *FOXA2* and *KLF5*) [26, 27, 28, 29, 30]. Among these, *SOX9* encodes a transcription factor controlling intestinal crypt homeostasis by blocking intestinal differentiation and promoting an intestinal stem cell-like program, and *SOX9* mutations have been reported in 29% of genome stable colorectal cancers (CRC) [30, 31]. We noted a higher prevalence of *SOX9* mutations in GCEP compared to TCGA GCs (Fisher test p-value 6.2x10^-8^). In GCEP, the majority of *SOX9* mutations were C-terminal truncating exon 3 mutations (truncating mutations: 119/173; 69%; truncating mutations at exon 3; 80/119; 67.2%) similar to CRC (**Figure 1D**) and were more common in antral biopsies compared to body/cardia samples (18.4% vs 5.1%; Fisher-test p-value 6.9x10^-12^). Mining TCGA expression data, we found that *SOX9* C-terminal truncating mutations were significantly associated with higher *SOX9* RNA expression in both GC and CRC cohorts (GC : log2 fold change 0.86; adjusted p-value 9.8x10^-3^; CRC: log2 fold change 0.73; adjusted p-value 6.2x10^-10^) (**Figure 1E**), supporting previous reports that *SOX9* truncating exon 3 mutations are associated with higher SOX9 protein expression [32]. In two independent GC datasets, high *SOX9* expression was observed in CIN GCs, a molecular subtype associated with IM (Wilcoxon test p-value 1.6x10^-7^ for TCGA and 4.9x10^-6^ for GASCAD) (Supplementary Figure 4). Increased *SOX9* RNA expression was significantly correlated with stemness scores in GC (TCGA GC: Spearman rho 0.35, p-value 1.5x10^-10^, GASCAD: Spearman rho 0.48, p- value 2.1x10^-8^), and *SOX9* mutated GCs showed expression signatures of oxidative phosphorylation (Normalized enrichment score, NES 2.6; adjusted p-value 3.6x10^-16^) and MYC pathway targets (NES 2.8; adjusted p-value 3.2x10^-20^) (**Figure 1F, G**). These findings suggest that *SOX9* mutations in IM may promote intestinal stem cell lineages and clonal expansion. However, the reduced frequency of *SOX9* mutations in GC suggests that *SOX9-*mutated IM clones may not be obligate precursors of malignancy.

### Spatiotemporal Clonal Dynamics in Normal, IM, and Dysplastic Gastric Tissues

Expansions of genetically-related cell populations (“clones”) in histologically normal tissues may pose a risk factor for malignancy [20, 33]. To ask if clone sizes differ between different categories of gastric pre-malignancy, we determined clone sizes as twice the mutation VAF [34] and estimated the fractional size of a gastric tissue covered by mutant drivers from the total summed size of driver clones in each biopsy (capped at 1.0) [34]. We found that biopsies from IM subjects were often polyclonal (median clone size 3.2%) while similar-sized clones were rare in normal subjects (median size 0%; Wilcoxon test p-value 1.9x10^-14^). Clone sizes expanded further in biopsies concurrent with dysplasia particularly in the antrum (median size 13.2%; p-value 4.3x10^-4^) but not body/cardia (median size 1.4%; p-value 0.50) (**Figure 2A**). The latter finding is consistent with GCEP clinical observations, where the majority of dysplastic and early GC lesions emerged from the antrum.

**Figure 2.**
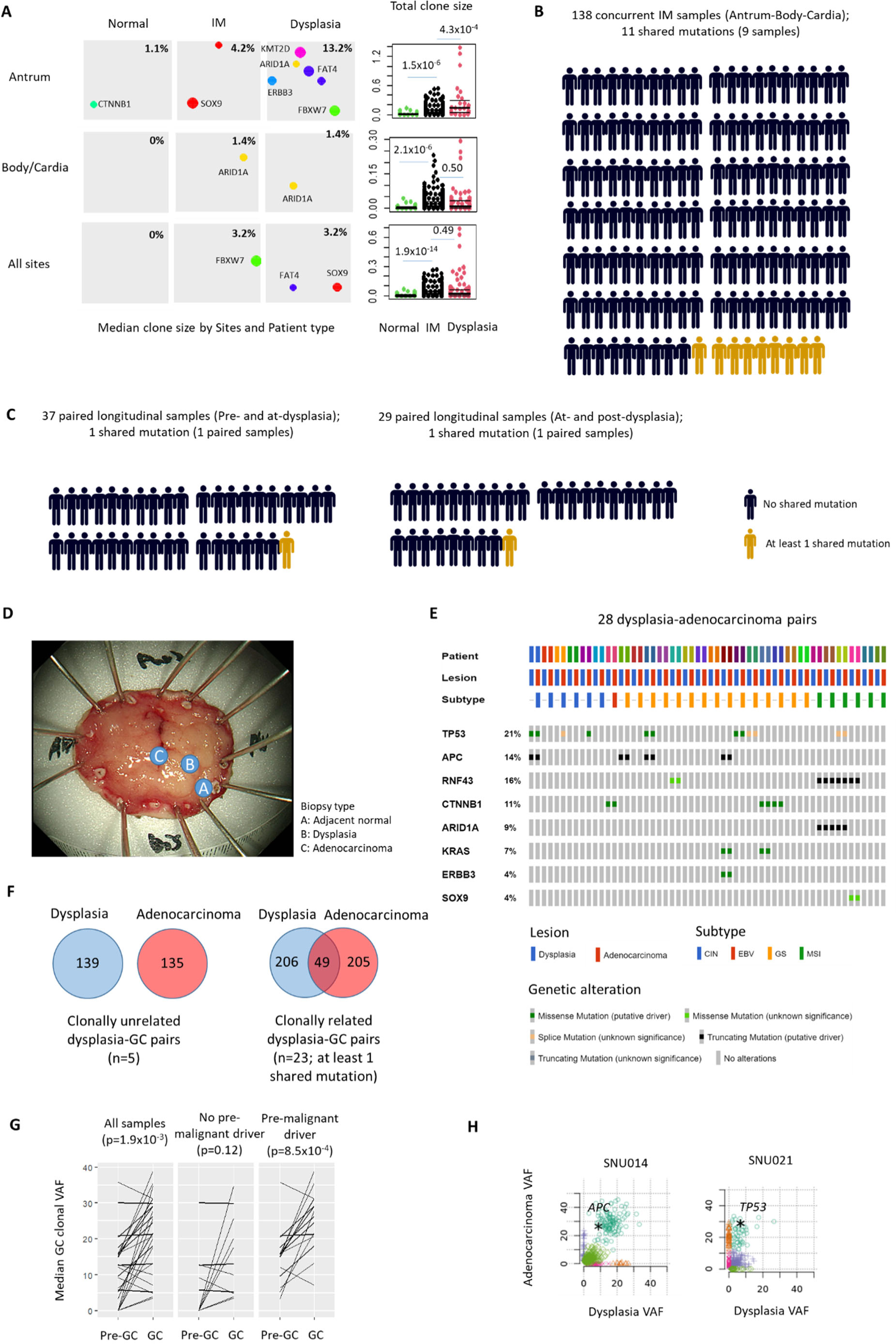
Clonal dynamics in IM, dysplasia and early GC. (A) Bubble plots showing predicted genetic clones in representative normal, IM and dysplasia samples. Sizes of driver clones were inferred from VAF values observed in various sample types. Beeswarm plots shows the total size of clones in all normal, IM and dysplasia samples by stomach region or across all regions. (B) Shared (gold) and private (black) somatic mutations observed in pre-malignant samples sampled from different stomach sites in the same subject (n=138). (C) Shared (gold) and private (black) somatic mutations observed in longitudinal samples from the same subject, either (left) from pre-dysplasia to dysplasia (n=37) or dysplasia to post-dysplasia (n=29). (D) WES on samples exhibiting concurrent normal, dysplasia and regions of early GC. (E) Oncoplot showing selected GC driver genes in 28 dysplasia-early GC pairs. Many mutations observed in dysplasia are also observed in regions of concurrent GC. (F) Sharing of mutations in clonally related (n=23) and unrelated (n=5) dysplastic-GC pairs. Median numbers of shared and private mutations in dysplasia and GC lesions are indicated. (G) Median clone sizes in dysplastic and GC samples, with or without identified driver mutations in the dysplastic lesion. (H) SciClone 2D plot showing clonal expansions associated with selected driver genes (*APC*, *TP53*) in dyplasia and concurrent GC.

To ask if clones are shared between IMs from different stomach regions in the same subject (“intra-subject”), we analysed 115 IM subjects where multiple biopsies from different stomach regions (antrum, body and cardia) were sampled at the same time point (138 antral/body/cardia trios in total). Only 8 subjects (9 samples) had IMs from different regions sharing at least one mutation, with the vast majority of subjects exhibiting genetically unrelated clones (**Figure 2B**). Further, to ask if these clones are stable or fluctuate dynamically over time, we then analysed 66 matched longitudinal pairs from the same subject, where IMs were sampled at different time points (37 pairs: at pre-dysplasia and adjacent to dysplasia; 29 pairs: at adjacent to dysplasia and subsequent regression of dysplasia). Shared mutations were observed in only 2 subjects (3.0%), suggesting that most IM clones are highly dynamic and transient (**Figure 2C**).

We hypothesized that in contrast to IM where clones are transient, clones in dysplastic gastric tissues might be more persistent contributing to their larger sizes. To explore this possibility, we applied WES to 28 GC patients from South Korea, where in each patient normal gastric tissue, dysplastic tissue, and early GCs were concurrently sampled (**Figure 2D**). In the matched GC-dysplasia pairs, the majority of driver gene mutations (22/26) observed in GC were also observed in the patient-matched dysplasia (*TP53*, *APC*, *ARID1A*, *RNF43*, *KRAS*, *ERBB3*, *CTNNB1*, *SOX9*) (16/20 GCs; 8 GCs had no identifiable driver mutations) (**Figure 2E**), with most pairs (23/28) showing at least one shared mutation between dysplastic lesions and matched GCs (**Figure 2F**). Clonal reconstructions using SciClone predicted clonal expansions from dysplastic (median clonal VAF 12.4%) to malignant GC lesions (median clonal VAF 21.4%) (paired Wilcoxon test p-value 1.9x10^-4^; 26 pairs), with more pronounced expansions in dysplastic lesions containing driver mutations (n=15; paired Wilcoxon test, p-value 8.5x10^-4^) (**Figure 2G**, **H**), consistent with the latter driving clonal expansion into malignancy. These spatiotemporal results suggest that in IM, independent clones can arise at different stomach sites, but the majority of these IM clones are likely transient which may be caused by high turnover rates. In contrast, genetic clones in dysplastic tissues may be more persistent, increasing the likelihood of transiting to full malignancy.

### IM scRNA-seq reveals shifts in gastric tissue ecology with expansions of intestinal cell lineages

We sought to define the repertoire of intestinal lineages in IM and their interplay with gastric lineages. We performed scRNA-seq (single-cell RNA sequencing) of antral IMs from 10 non-cancer subjects exhibiting different levels of IM severity (6 negative/mild; 3 moderate; 1 severe). After excluding low quality and doublet cells, we performed Leiden clustering on 42,570 cells and identified 24 cell clusters belonging to 4 major cellular lineages, including gastric (36.7% of cells; marked by *TFF2*), intestinal (27.3%; *REG4*), immune (21.8%; *SRGN*), and stromal cells (7.6%; *DCN*) (**Figure 3A**, Supplementary Figure S5A). Focusing on the gastric and intestinal lineages, we identified four gastric lineages based on previously reported marker genes, including gastric stem cells (*IQGAP3, STMN1, MKI67*), isthmus cells (*SULT1C2, CAPN8, TFF1*), LYZ-positive cells (*LYZ, MUC6, PGC*; which are mucous-secreting cells at the lower part of gastric glands [35, 36, 37]) and immature and mature pit cells (*GKN1, GKN2, TFF2*) (Supplementary Figure S5B). Similarly, we identified four intestinal-type lineages, including intestinal stem cells (*OLFM4, CDCA7*), transit amplifying cells (*DMBT1*), enterocytes (*FABP1, FABP2, KRT20*) and goblet cells (*SPINK4, MUC2, TFF3*) (Supplementary Figure S5C). Consistent with histology, IM severity correlated with increased proportions of intestinal lineages (Pearson correlation, r=0.82, p-value 3.8x10^-3^), including intestinal stem cell (r=0.63), transit amplifying cells (r=0.54) and enterocytes (r=0.69), and decreases of gastric lineages (r -0.79, p-value 7.0x10^-3^), including LYZ-positive cells (r=-0.18) and isthmus cells (r=-0.75) (**Figure 3B**). No significant differences were observed in the proportions of immune (Pearson correlation -0.28, p-value 0.46) or stromal cells (Pearson correlation -0.37, p-value 0.32).

**Figure 3.**
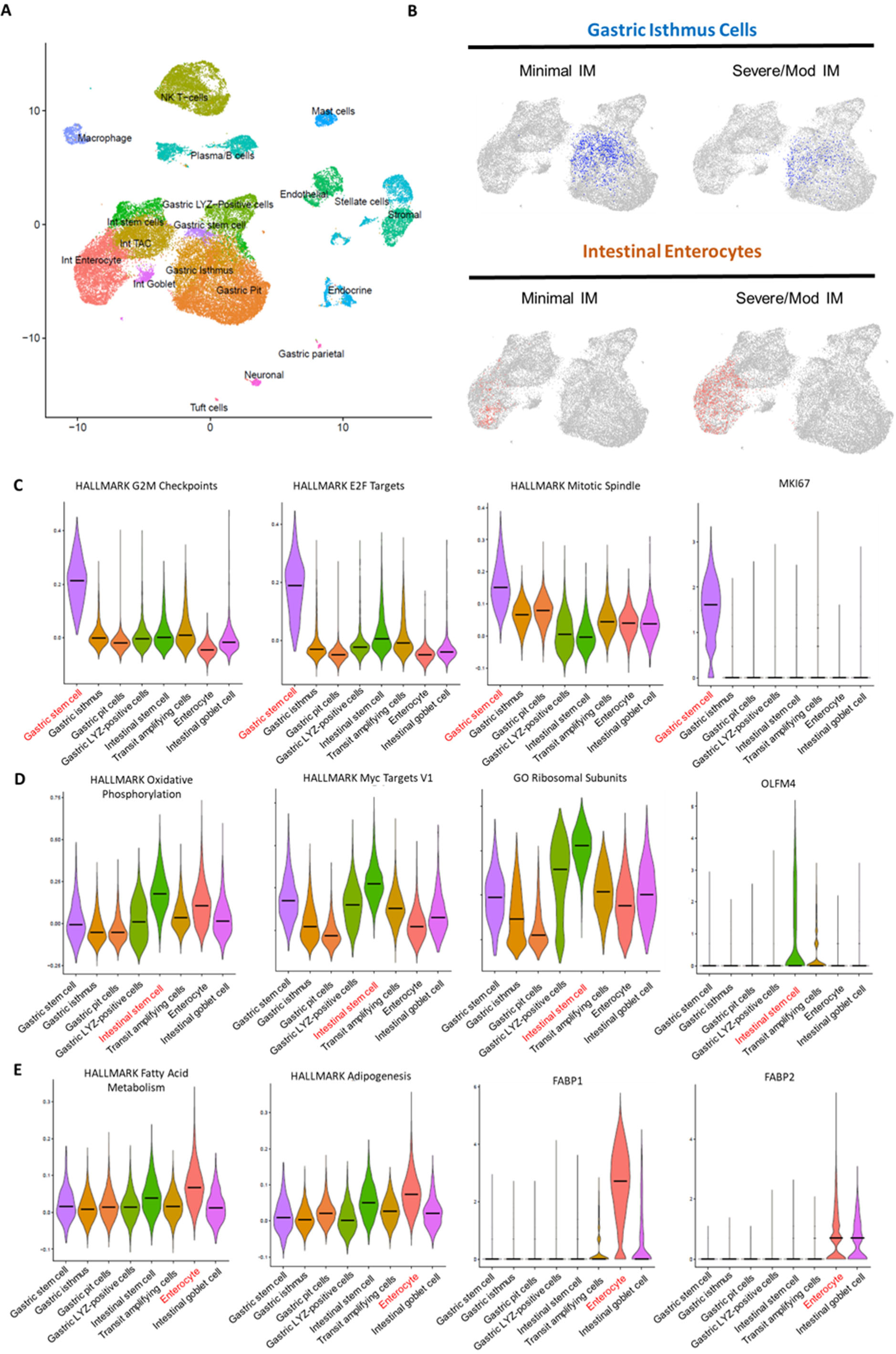
Single cell transcriptomic landscape of IM. (A) 24 cell types/lineages identified from single-cell RNAseq profiling of antral IMs. (B) Increased proportions of intestinal lineages (enterocyte, brown) cells and decreased gastric lineage cells (gastric isthmus, blue) in subjects with severe/moderate IM compared with subjects with mild/negative IM. (C) Violin plots showing enrichment of cell cycle pathways in gastric stem cell lineages. (D) Violin plots of oxidative phosphorylation and Myc target V1 pathways reveals highlights expression in intestinal stem cell lineages. Also shown are expression levels of the intestinal stem cell marker *OLFM4*. (E) Violin plots showing enrichment of fatty acid metabolism and adipogenesis pathways in intestinal enterocyte lineages. Intestinal enterocytes are marked by expression of *FABP1* and *FABP2*.

We proceeded to dissect functional pathways associated with the gastric and intestinal lineages. Gastric stem cells marked by *IQGAP3* up-regulated pathways related to cell division including G2M checkpoint (NES 3.4, adjusted p-value 2.5x10^-36^), E2F targets (NES 3.3, adjusted p-value 7.6x10^-31^), mitotic spindle (NES 2.9, adjusted p-value 1.6x10^-15^), and higher expression of the proliferation-related genes *MKI67* (93% vs 15.5%; adjusted p-value 2.1x10^-280^) and *TOP2A* (92% vs 13.2%; adjusted p-value 3.6x10^-298^) (**Figure 3C**). While normally quiescent, the presence of highly proliferative gastric stem cells may reflect ongoing cellular regeneration in response to tissue injury caused by HP infection, which is required for IM development [38]. Intestinal stem cells marked by *OLFM4* were highly enriched in oxidative phosphorylation (Normalized enrichment score, NES 4.0; adjusted p-value 2.2x10^-31^) and MYC target V1 pathways (NES 3.8; adjusted p-value 1.2x10^-26^), consistent with adult stem cells switching to mitochondrial oxidative phosphorylation when transitioning to a more proliferative state, along with up-regulation of ribosomal genes [39, 40] (**Figure 3D**). Indeed, small (RPS, n=30) and large (RPL, n=45) ribosomal subunits encompassed 75% of the top 100 up-regulated (by fold change) genes in the intestinal stem cell cluster. Compared to intestinal stem cells, intestinal enterocytes (which are more differentiated) marked by *FABP1*/*2* exhibited up- regulation of oxidative phosphorylation to a lesser degree (NES 2.4; adjusted p-value 1.6x10^-7^) and down-regulation of MYC pathways (NES -2.6, adjusted p-value 2.1x10^-6^) along with high expression of adipogenesis (NES 2.3; adjusted p-value 5.0x10^-5^) and fatty acid metabolism programs (NES 1.9; adjusted p-value 5.5x10^-3^) (**Figure 3E**). Notably, *SOX9* was highly expressed in gastric LYZ-positive cells (49.7%, adjusted p-value < 1.0x10^-300^), intestinal stem cells (27.6%, adjusted p-value < 1.2x10^-42^) and transit amplifying cells (27.5%, adjusted p-value < 9.6x10^-21^) with intestinal stem cells expressing high levels of several *SOX9*-associated expression signatures (eg stemness, oxphos, MYC targets). This finding suggests that the presence of the latter signatures in bulk transcriptome data (see **Figure 1**) is likely linked to the increased proportion of intestinal stem cells rather than a uniform increase of *SOX9* signature expression across all cell types.

### Intestinal Stem-cell Dominant IM Exhibits Transcriptional Similarities to GC

To investigate relationships between the gastric and intestinal lineage heterogeneities observed in IM with malignant GC, we then integrated the IM scRNA-seq data with previously published scRNA-seq data from early-stage GCs (for this analysis, GC scRNA-data was restricted to epithelial cells exhibiting inferred somatic CNAs) [41] (Supplementary Figure 6A). Overall clustering of the combined IM and GC data confirmed close similarities between IM and GC epithelial cell populations (**Figure 4A**). Pseudotime analysis revealed two separate developmental lineage roots – one reflective of normal gastric lineages and another marked by intestinal lineages. Monocle3 trajectory analysis projected that early GC cells appear to be most closely related to intestinal stem-cell lineages, and more distantly related to other intestinal-related lineages such as differentiated enterocytes or goblet cells (**Figure 4B**). These findings may suggest that intestinal stem-cell subpopulations in IM may harbour a potential cellular reservoir for the emergence of intestinal-type GC. Indeed, some *OLFM4*-expressing intestinal stem-cells also co-expressed *LGR5* (147/855 cells; 17.2%; Fisher test odds ratio 22.4, p-value < 2.2x10^-16^) and *AQP5* (227/855 cells; 26.5%; Fisher test odds ratio 7.7, p-value < 2.2x10^-16^), which are both gastric stem cell markers previously proposed to mark cancer stem cells [42].

**Figure 4.**
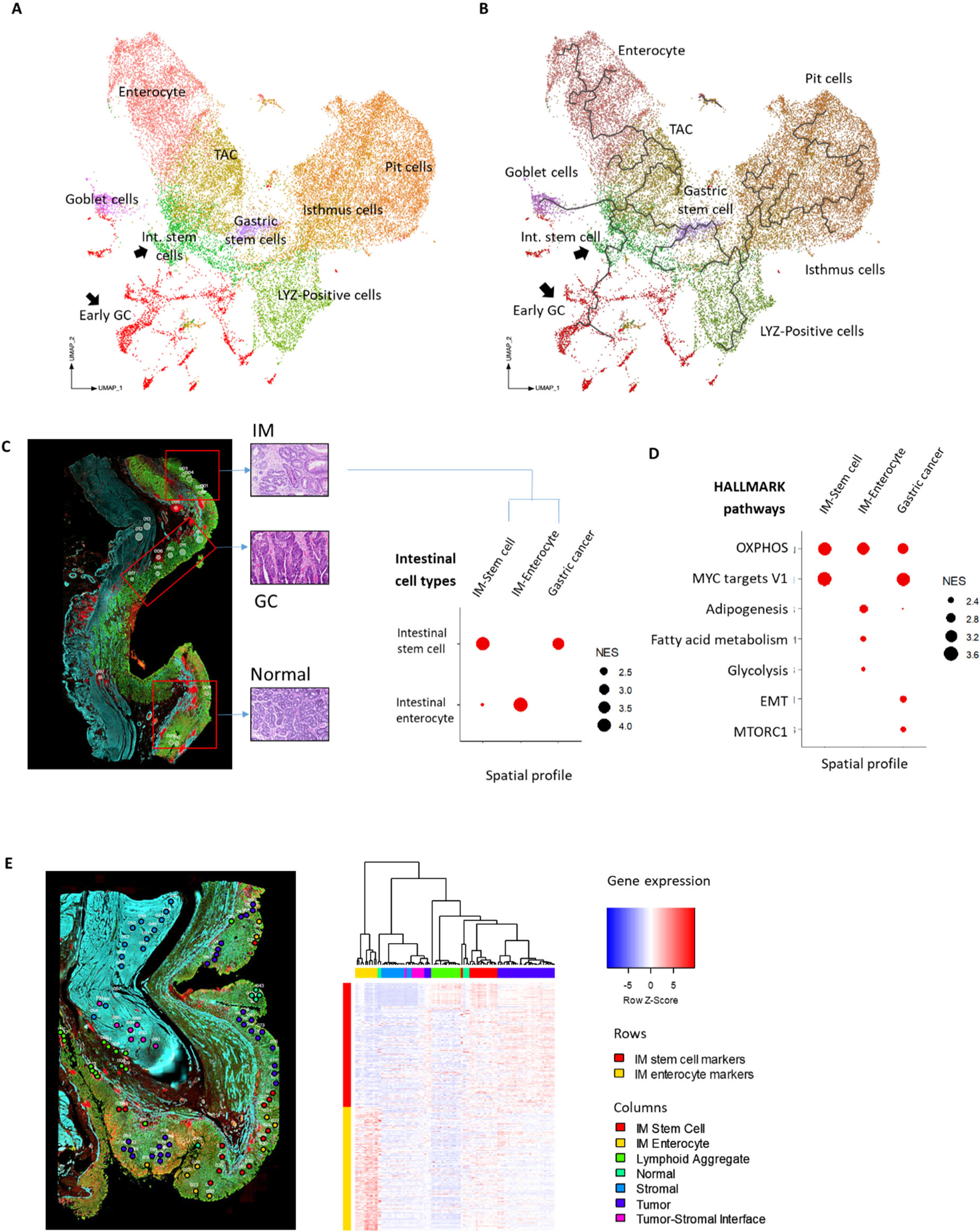
Trajectory analysis of IM and GC cells. (A) UMAP plot showing the clustering of single cells from IM patients and early GC cells. Early GC scRNA-seq profiles were obtained from [41]. GC cells and intestinal stem cells are marked by black arrows. (B) Monocle3 trajectory analysis. GC cells are most closely related to intestinal stem cells. (C) Representative AOIs from a tissue section displaying concurrent normal, IM and GC (left). AOIs/ROIs from IMs were annotated as stem-cells dominant IM (IM-stem cell) or enterocyte dominant (IM-Enterocyte) based on scRNA-seq expression profiles (right). (D) Dotplots showing enrichment of selected HALLMARK pathways in intestinal stem cell dominant IM, enterocyte-dominant IM, and GC. GCs are observed to also exhibit signatures of EMT and MTORC1 (E) Image of histological slide labelled with selected ROIs (left). IM regions were annotated as intestinal stem cell-dominant or enterocyte-dominant IM. Hierarchical clustering using IM stem cell and enterocyte markers of selected ROIs shows similarities between GC spatial profiles and intestinal stem-cell dominant IM (right).

To orthogonally confirm that intestinal stem-cell lineages in IM are related to GC, we then performed spatial transcriptomics using Nanostring Digital Spatial Profiling on tissue sections from 8 GC patients harbouring concurrent normal, IM and GC regions. Across 87 IM AOIs/ROIs, we calculated enrichment scores to annotate each IM region as “stem-cell dominant (n=37)” or “enterocyte-dominant (n=30)” using expression signatures from the scRNA-seq data (**Figure 4C**). Interestingly, we observed a significant negative correlation between HALLMARK inflammation scores with stem-cell dominant IM scores (rho -0.45, p-value 7.0x10^-4^), and a positive correlation with enterocyte-dominant IM scores (rho 0.63, p-value 6.8x10^-7^) (Supplementary Figure S6B), suggesting that IM stem cells occupy an immune-excluded niche. Consistent with this possibility, we observed a significant enrichment of IM enterocyte scores in CD45+ (n=6) compartments compared to CD45- regions (n=7) (Enrichment score 0.73 vs 0.38, Wilcoxon test p-value 1.2x10^-3^) (Supplementary Figure S6C).

Biological pathways activated in stem cell-dominant IM, enterocyte-dominant IM and GC were inferred using HALLMARK [43]. Consistent with the scRNA-seq data, stem-cell dominant IMs overexpressed oxidative phosphorylation gene sets (NES 3.5, adjusted p-value 1.4x10^-40^) and MYC targets V1 pathways (NES 3.7, adjusted p-value 4.2x10^-49^) which were notably also expressed in GC regions (OxPhos - NES 3.1, adjusted p-value 1.1x10^-27^; MYC - NES 3.5, adjusted p-value 2.2x10^-44^) (**Figure 4D**). Reciprocally, GC regions showed enrichment of gene expression programs related to intestinal stem-cells (NES 3.5, adjusted p-value 4.7x10^-78^) rather than differentiated enterocytes (NES 1.5, adjusted p-value 1.7x10^-4^). Pathways specific to enterocyte-dominant IM included fatty acid metabolism (NES 2.4, adjusted p-value 1.1x10^-7^), and adipogenesis (NES 2.7, adjusted p-value 1.2x10^-12^) which were not strongly up-regulated in GC regions (Fatty acid metabolism - NES 1.5, adjusted p-value 0.038; adipogenesis - NES 2.1, adjusted p- value 5.3x10^-7^). Compared to both stem-cell dominant and enterocyte-dominant IM, GC regions harboured additional signatures not observed in IMs such as genesets associated with epithelial-mesenchymal transition (NES 2.4, adjusted p-value 4.6x10^-11^) and MTORC1 signalling (NES 2.3, adjusted p-value 2.2x10^-9^). As an illustration, we performed hierarchical clustering on the spatial transcriptomics data from a single slide (93 ROIs). Hierarchical clustering using markers of intestinal stem-cells and enterocytes, grouped stem cell-dominant IMs together with GC while enterocyte-dominant IMs were more distantly related (**Figure 4E**). These results demonstrate that even in the same subject, IMs display significant lineage heterogeneity with stem-cell dominant IM exhibiting expression signatures similar to malignant cell populations.

### Bulk transcriptome sequencing across subjects identifies distinct expression subtypes of IM

Previous studies have underscored the biological and clinical relevance of expression-based molecular subtypes in cancer [44, 45]. To ask if IMs can be classified into distinct categories based on mRNA profiles, we then analyzed bulk RNA-seq transcriptomes across 183 pre-malignant GC samples including antrum (24 normal, 31 IM) and body/cardia (22 normal, 106 IM) samples. Initial expression based clustering of the normal gastric samples confirmed a distinct separation of antral and body/cardia samples, consistent with each stomach anatomical region being histologically distinct (**Figure 5A**). Using the normal antral and body/cardia dichotomy as a foundation, we then overlaid unsupervised hierarchical clustering of the IM gene expression data, revealing three distinct IM subtypes. The first IM subtype comprised antral IMs with expression similarities to antral gastric tissues (28/31), and the second subtype comprised body/cardia IMs with expression similarities to body/cardia normal tissues (65/106). However, we noted a third subtype comprising IMs from the stomach body/cardia but expressing transcriptional similarities with antral IMs (41/106) (**Figure 5B**). This phenomenon is reminiscent of ‘pseudo-antralization’, a process associated with HP infection, IM, and GC characterized by the appearance of antral-type mucosa in the body/cardia [46]. In keeping with this nomenclature, we hereafter refer to this third IM subtype as ‘pseudo-antralized IMs’.

**Figure 5.**
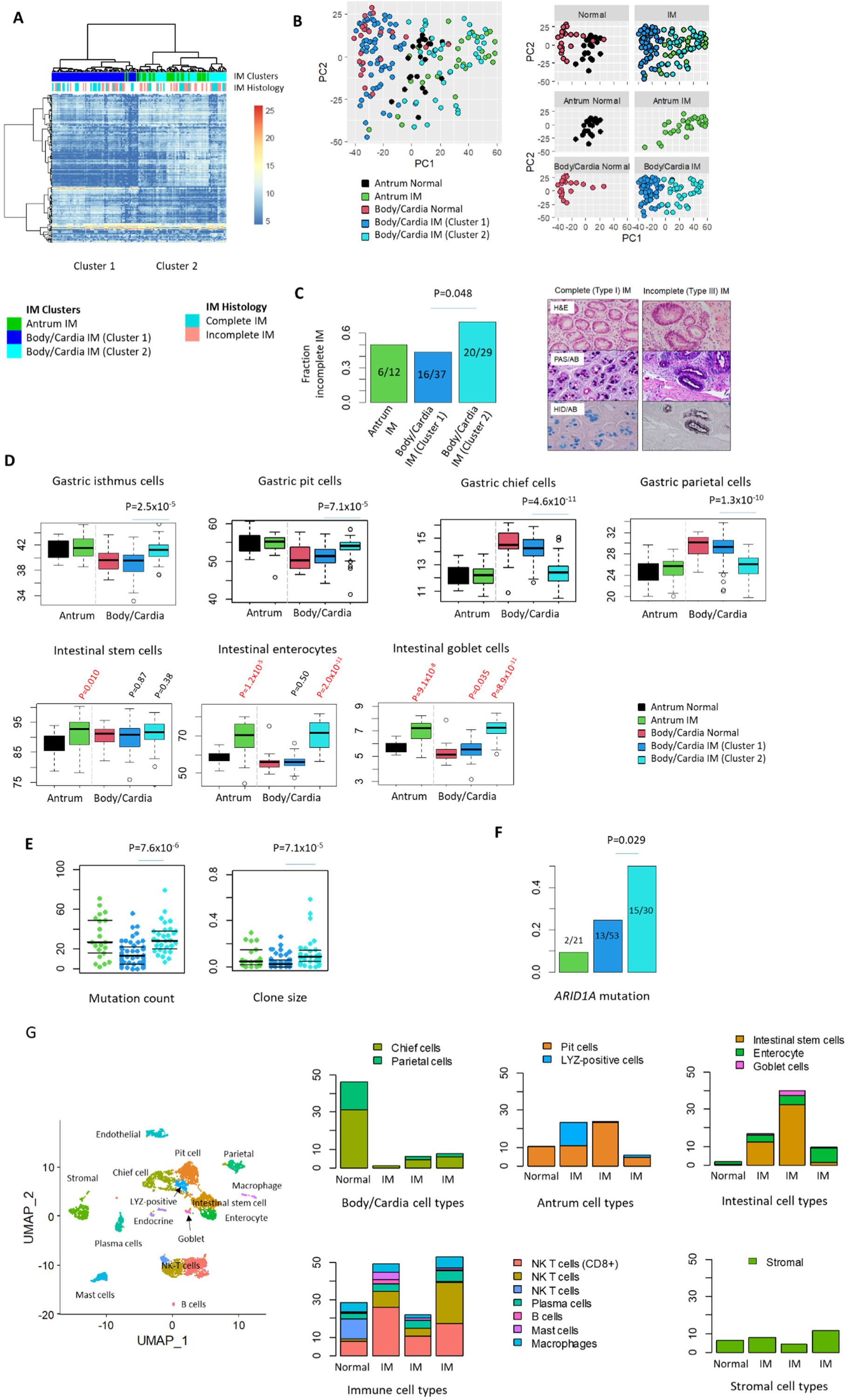
Expression-based Molecular Subtypes of IM and Pseudoantralization. (A) Hierarchical clustering of bulk IM RNAseq transcriptomes (n=137 IM). A cluster of body/cardia IMs (cluster 2, light blue) cluster with antral IMs (green). (B) PCA graphs of normal gastric samples and IMs. Normal antral and body/cardia samples were well demarcated, while IM samples are distributed across both regions. IM cluster 2 samples cluster with antral IMs. (C) Fraction of histologically-defined incomplete and complete IM subtypes across IM expression subtypes (left). Representative images of Type I complete and Type III incomplete IM (right; adapted from Huang et al. 2018). (D) ssGSEA scores for gastric cell types and intestinal cell types in antral and body/cardia normal samples and IMs. Cluster 2 IMs exhibit similarities to antral IMs. (E) Mutation counts and clone sizes of IM expression subtypes. Cluster 2 body/cardia IMs exhibit higher mutation counts and clone sizes relative to Cluster 1 body/cardia IMs. (F) *ARID1A* mutations are enriched in Cluster 2 body/cardia IMs. (G) Proportion of antral, body/cardia, intestinal, immune and stromal cell types from scRNA- seq of gastric body biopsies (n=4).

Several lines of evidence support pseudo-antralized IMs as a distinct molecular entity. First, when correlated to histology, pseudo-antralized IMs were significantly associated with incomplete IM histology (containing mixtures of goblet, enterocyte and immature mucosal cells) (**Figure 5C**; Fisher-test p-value 0.048), a histological subtype associated with higher GC risk [47]. Second, compared to body/cardia IMs, pseudo-antralized IM harboured increased gene expression programs of antral cell types (gastric pit; Wilcoxon test p-value 7.1x10^-5^ and isthmus cells; p-value 2.5x10^-5^), and mature intestinal cell lineages (enterocyte; p-values 2.0x10^-11^ and goblet cells; p-values 8.9x10^-11^), with reduced expression of body/cardia cell types (gastric chief; p-value 4.6x10^-11^ and parietal cells; p-value 1.3x10^-10^) (**Figure 5D**). Third, across the subset of GCEP samples (104 cases) with both DNA mutation and RNAseq data, pseudo-antralized IMs exhibited significantly higher mutation rates (Wilcoxon test p-value 7.6x10^-6^) and clone sizes (p-value 7.1x10^-5^) compared to body/cardia IMs and similar to antral IMs (**Figure 5E**). Fourth, pseudo-antralized IMs exhibited a higher frequency of *ARID1A* mutations compared to body/cardia (Fisher-test p-value 0.029) or antral IMs (Fisher test p-value 0.0028) (**Figure 5F**). Taken collectively, these observations suggest that pseudo-antralized IMs, while resident in the body/cardia are distinct from body/cardia IMs, and while similar to antral IMs in many respects, are also distinct from native antral IMs by virtue of both anatomic location, higher presence of *ARID1A* mutations, and (as shown in the next section) a distinct microbial and inflammatory milieu.

Pseudo-antralized IMs exhibited features reminiscent of SPEM (see Discussion). To further validate the bulk RNA-seq results, we performed single-cell RNA sequencing on 4 gastric body biopsies (3 IMs and 1 normal; Supplementary Figure S7A). We identified 18 cell clusters corresponding to gastric body lineages (chief and parietal cells), gastric antral cells (LYZ-positive cells and pit/isthmus cells), intestinal lineage cells (intestinal stem cell and enterocytes) and immune cells (Supplementary Figure S7B). Supporting the accuracy of our anatomic sampling, the normal body biopsy contained a higher proportion of chief and parietal cells (46.4%) compared to normal antrum biopsies (average 0.24%), and a lower percentage of antral cell types (10.4% vs 45.6%). Consistent with ‘pseudo-antralization’, we further observed a depletion of normal body cell types (4.9% vs 46.4%) but an increase in normal antral (17.7% vs 10.4%) and intestinal cell types (22.2% vs 1.8%) in body IM biopsies (Figure 5G). In one body IM sample, we observed an increase in LYZ- expressing cells which are abundant in normal antrum but rare in the normal body. These cells also co-expressed *AQP5*, consistent with *AQP5* being a marker of SPEM [48].

### Pseudo-antralized IMs exhibit an inflammatory microenvironment associated with a distinctive oral microbial community

We observed a higher proportion of immune cell types in body IM biopsies compared to antrum IM (41.4% vs 25.6%; Wilcoxon test, p-value 0.27), suggesting that IM emergence in the gastric body may be associated with a specific immune microenvironment. Notably, while DNA-based alterations can capture changes only in epithelial cells, bulk RNA profiles can also provide insights into alterations affecting other non-epithelial cellular populations including immune cells. We found that pseudo-antralized IMs and body/cardia IMs exhibited increased TNFA signalling via NFKB (pseudo-antralized IM – NES 2.0, adjusted p-value 1.3x10^-5^; body/cardia IM - NES 2.3, adjusted p-value 4.7x10^-8^) (**Figure 6A**) suggesting that IMs present in the body/cardia are associated with increased inflammation. In particular, pseudo- antralized IMs exhibited increased interferon alpha (NES 2.5; adjusted p-value 7.0x10^-11^) and interferon gamma responses (NES 2.6; adjusted p-value 1.5x10^-14^) exceeding that observed in native body/cardia IMs. Using two different cell deconvolution tools (CIBERSORTX and ESTIMATE; **Figure 6B**), we confirmed significant increases of immune cells in pseudo-antralized IMs (Wilcoxon test p-value 1.3x10^-5^ in pseudo-antralized IM). Interestingly, these immune cell changes were largely associated with increases in memory B cells (Wilcoxon test p-value 8.0x10^-5^) and a corresponding decrease in CD8 T cells (Wilcoxon test p-value 8.0x10^-4^).

**Figure 6.**
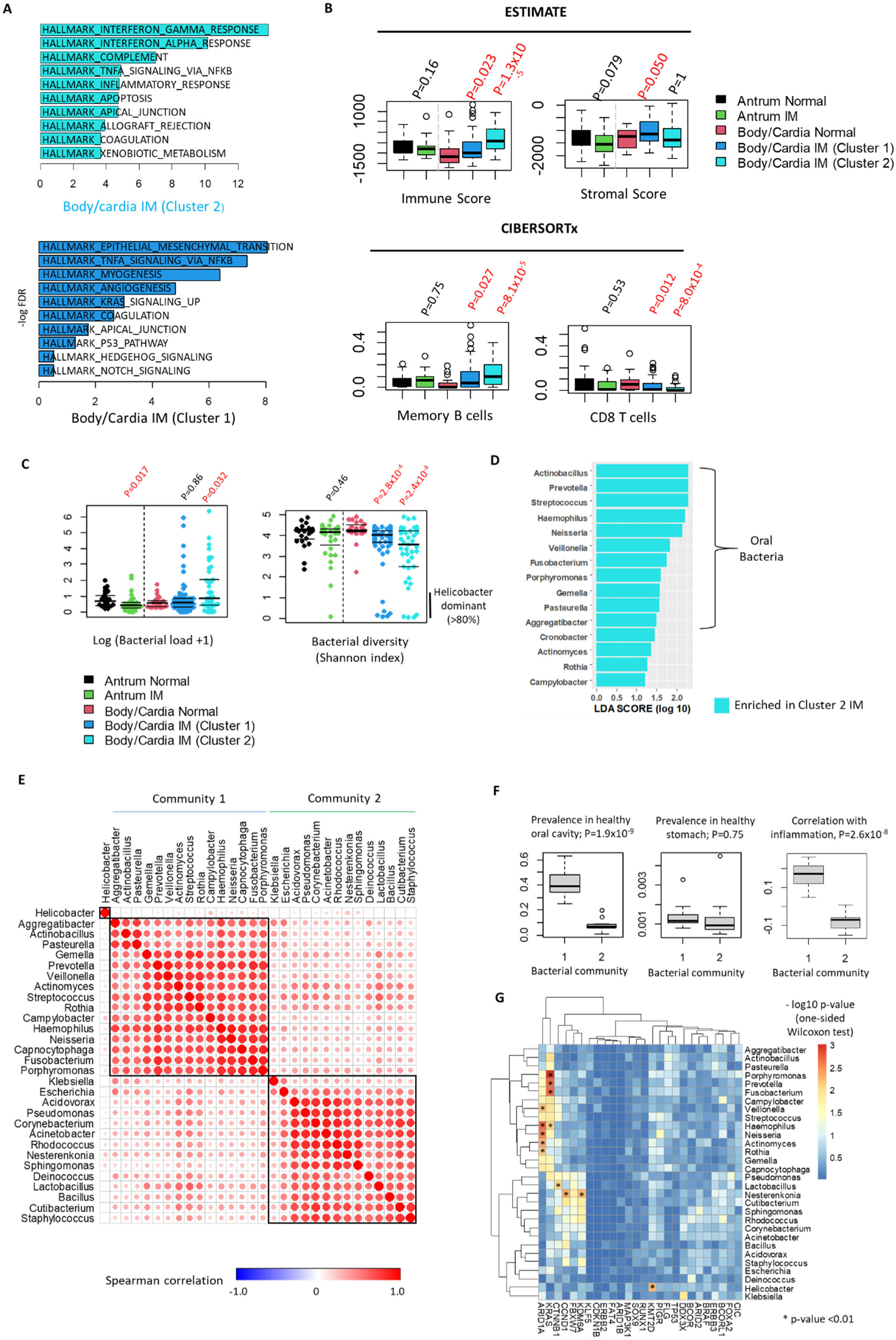
Immune landscape in IM. (A) GSEA of expression signatures in body/cardia IM subtypes 1 and 2. Inflammatory signatures (Interferon gamma, etc) are upregulated in Subtype 2. (B) Immune and stromal content deconvolution analysis using ESTIMATE and CIBERSORTx. Body/cardia subtype 2 samples exhibit upregulation of immune scores and B-cell programs. (C) Bacterial density and diversity in IM and normal samples. Body/cardia IM subtype 2 samples exhibit increased bacterial loads but lower diversity. (D) LDA analysis comparing microbial genus between body/cardia IM subtypes 1 and 2. (E) Correlation analysis (Spearman) of the 30 most abundant bacterial genus identified in this study. The 30 genera represent the major contributors to microbial levels in this study. Two distinct microbial communities are observed (C1 and C2). (F) Prevalence of bacterial genus from C1 and C2 in reference microbiomes from oral cavity (left) and normal stomach (middle). Correlation between community C1 with HALLMARK inflammation scores (right). (G) Association between bacterial genus abundance with somatic driver mutations in IM samples. Bacterial genera positively associated with somatic mutations are indicated with asterisks (p<0.01).

We hypothesized that the inflammatory environment observed in pseudo- antralized IMs might be caused by alterations in microbial composition. To investigate this possibility, we used Pathseq [49] to estimate bacterial content and diversity from the RNAseq data at the genus level. Compared to DNA-based measurements, inferring microbial identities based on RNA enables the identification of transcriptionally active bacterial communities rather than remnants of previous infection [50]. Of ∼34 million bacterial reads from 847 bacterial genera in the 183 samples, reads mapping to Hp accounted for 79.3% of all unambiguously mapped bacterial reads. *Helicobacter* RNA reads were enriched in IM subjects compared to normal subjects (Wilcoxon test, p-value 7.1x10^-3^). However, pseudo-antralized IMs exhibited both increased bacterial levels compared to body/cardia normal samples (Wilcoxon test p-value 0.032) and also reduced diversity (p-values 2.8x10^-4^ in non- antralized IM, 2.4x10^-4^ in pseudo-antralized IM) (**Figure 6C**). The coupling of increased bacterial load with decreased diversity (sometimes termed “microbial dysbiosis”) has been linked to various diseases such as rheumatoid arthritis [51] and diabetes [52].

We deepened our analysis to identify microbial communities specifically associated with inflammation in pseudo-antralized IM. Linear discriminant analysis (LDA) highlighted bacterial communities comprising *Streptococcus*, *Prevotella* and *Fusobacterium* in pseudo-antralized IM (LDA score 1 to 3) compared to non- antralized IM (**Figure 6D**). A more refined clustering analysis of the top 30 most abundant bacterial genus in the RNA-seq data yielded two clusters of bacterial communities (**Figure 6E**). Cluster 1 comprised bacteria normally associated with the oral cavity (e.g. *Streptococcus*, *Porphyromonas*) (Wilcoxon test p-value 1.9x10^-9^) but typically absent in healthy stomach (p-value 0.75) compared to cluster 2 (e.g. *Acidovorax*, *Pseudomonas*). These observations support previous studies employing 16S sequencing reporting that certain oral bacteria may be associated with IM onset after *H. pylori* eradication [53] – we confirmed that our cluster 1 community overlaps significantly with these previous reports (Fisher-test p-value 6.3x10^-3^). Notably, levels of microbial cluster 1 were significantly associated with increased inflammation scores (**Figure 6F**; p-value 2.6x10^-8^), rendering it possible that presence of these microbes may initiate a pro-inflammatory process. We also found that cluster 1 microbes were also more frequently associated with GCEP samples harbouring driver gene mutations such as *ARID1A* and *KRAS* (**Figure 6G**).

### Combined Genomic-Clinical Predictive Models Outperform Models Based on Clinical Information Only

Finally, to assess the clinical relevance of our molecular results, we evaluated if incorporation of genomic information might improve current clinical models used to stratify IM patients for dysplasia risk [19]. First, we focused on antral samples and used logistic regression analysis and ROC curves to benchmark the molecular variables against clinical features. We first compared the genomic features from antral samples at the time of dysplasia to the non-dysplasia subjects (**Figure 7A**). Univariate and multivariate analyses were performed for each risk factor. Multivariate logistic regression analysis showed that a positive pepsinogen index (B=1.768, 95%CI 0.519 to 3.017, p=0.006), smoking (B=1.363, 95%CI 0.249 to 2.477, p=0.016), higher mutation counts (B=0.04, 95%CI 0.005 to 0.075, p=0.023), and larger clone sizes (B=6.88, 95%CI 2.386 to 11.374, p=0.003) significantly increased the risk of dysplasia. Notably, integrated molecular and clinical models achieved superior performance as indicated by higher AUCs in predicting dysplasia (AUC=0.846, 95% CI 0.753 to 0.939, p<0.001) compared to clinical models alone (AUC=0.707, 95%CI 0.576 to 0.838, p=0.002).

**Figure 7.**
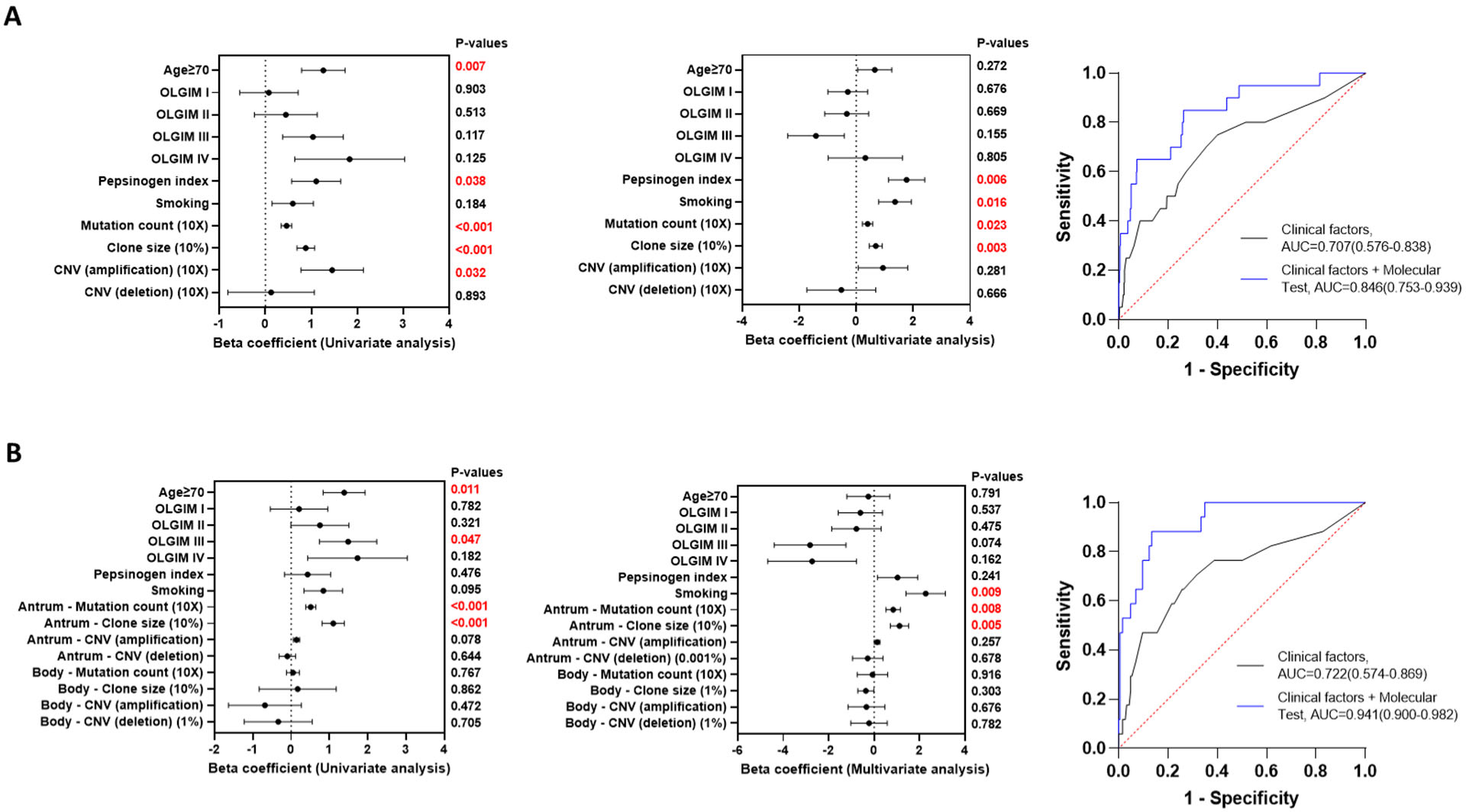
Predicting IM Progression Risk from Clinical and Genomic Features. (A) Clinical factors (age≥70, OLGIM score, pepsinogen index, smoking status) and genomic features (mutation count, clone size, copy number variation (CNA; amplification/ deletion) were used to stratify the risk of gastric dysplasia in patients with antral biopsies (Dysplasia n=23 vs Non-dysplasia n=599). Features were tested in both univariate and multivariate analysis. (Right) AUC curves showing accuracy of prediction based on clinical factors only (grey) or clinical and genomic factors (blue) (B) Analysis of patients with both antral and body biopsies (Dysplasia n=20 vs Non-dysplasia n=186). Left panel shows the forest plots of univariate and multivariate logistic regression analysis. The right panel shows ROC curves and corresponding AUC values to evaluate model performance.

As IM in the stomach body may represent a more advanced pathology, we further interrogated the cohort with molecular test results from both the antrum and body at the time of dysplasia (**Figure 7B**). Similar to the findings above, smoking (B=2.274, 95%CI 0.575 to 3.974, p=0.009), higher mutation count in the antrum (B=0.084, 95%CI 0.022 to 0.146, p=0.008), and larger clone size in the antrum (B=11.195, 95%CI 3.331 to 19.058, p=0.005) significantly increased the risk of dysplasia. The prediction accuracy of the integrated molecular and clinical model (AUC=0.941, 95%CI 0.9 to 0.982, p<0.001) was higher compared to clinical models (AUC=0.722, 95%CI 0.574 to 0.869, p=0.003). These observations suggest that integrating molecular information with clinical data is likely to improve prediction models to stratify the risk of subjects with gastric pre-malignancy.

### Discussion

To our knowledge, the present study reports the largest genomic and transcriptional survey of human IMs to date. Similar to GCs, IMs can involve different stomach regions, with IMs tending to originate in the antrum due to Hp infection [54]. Hp preferentially colonizes antral cell types such as pit cells [55] causing mucosal atrophy and IM [38, 56]. As atrophy/IM progresses, Hp levels often decrease due to IM cells being less hospitable to infection [57], raising the possibility that IM may function a protective mechanism against Hp. Hp may consequently disappear from the antrum but persist at other stomach regions [58] - in GC patients, Hp detection rates are thus often higher in the body due to lower atrophy and IM levels in the latter [58, 59].

Our results support a growing body of literature that IMs are not a homogenous entity but highly heterogenous between and even within patients. Not all IM patients will progress to GC, and histologically IMs can be classified into complete or incomplete subtypes (see Introduction) [60]. A meta-analysis of >407,000 subjects reported that incomplete IMs (pooled OR 9.48) were significantly associated with GC compared to complete IMs (pooled OR 1.55) [15]. GC onset was also higher among patients with IM involving the antrum and body (extensive IM; pooled OR = 7.39) compared to the antrum only (pooled OR = 4.06) [15]. These differences may be contributed at least in part by region-specific cellular populations in the stomach including stem cells. For example, antral isthmus stem cells are a potential stem cell population with high proliferative potential [61], and LGR5/AQP5- expressing stem cells in the antral gland base have also been identified as a potential source of IM and GC [42, 62]. In the gastric body, differentiated chief cells may contribute to GC by acting as reserve stem cells after epithelial injury [63], and lineage tracing in mice has revealed that chief cells can undergo transdifferentiation into SPEM [63] which is as strongly associated with GC as IM [64].

Recent sequencing advances have enabled the detection of somatic mutations associated with genetic clones (genetically identical subpopulations of cells) and subclones in normal, inflamed, and pre-malignant tissues [65]. In tissues such as the esophagus [33, 66] microscopic clones harboring driver mutations such as *NOTCH1* may eventually expand to macroscopic levels, with 50% of esophageal epithelium eventually colonized by mutant clones [33, 66]. In our study, *SOX9* was identified as a new IM driver gene in certain clones. In genome-stable CRC, *SOX9* is mutated in 29% of cases [31] with most *SOX9* alterations being nonsense/frameshift mutations preferentially clustering within the C-terminus [31] and leading to *SOX9* overexpression [32]. In CRC lines, *SOX9* silencing caused proliferation defects, while *SOX9* overexpression led to reduced expression of differentiation markers consistent with *SOX9* blocking intestinal differentiation in human CRC. The overlap of *SOX9* mutational profiles between CRC and IM suggests that *SOX9* mutations may also play an initiating role in IM, by impeding differentiation and promoting lineage transformations and stem-like states. However, while *SOX9* may promote IM clonal expansion, the lower frequency of *SOX9* mutations in GC suggests that not all *SOX9*-expanded IM clones may lead to cancer, similar to *NOTCH1* in esophageal cancer [67]. One possible explanation might be that IM clones are dynamic and transient, in contrast to dysplastic clones that are larger and more stable with a higher propensity to transmit oncogenic genetic alterations to eventual GCs. It is also possible that certain genes can act as drivers in non-malignant tissues but protect against subsequent cancer, as has been proposed for inflamed colonic tissues harboring clones mutated in genes such as *PIGR*, *NFKBIZ* and *ZC3H12A* [21, 22, 68] which all exhibit low mutation rates in CRC.

Our study reinforces an important role for metaplasia in cancer development where metaplastic cells co-expressing aberrant markers of multiple lineages have higher phenotypic plasticity and cancer propensity. Complementing bulk analysis, single-cell approaches are providing important insights into the cellular programs of metaplastic cells in the esophagus [69], stomach antrum [36] and colon [70]. These studies have shown that Barrett’s esophagus (BE) may originate from normal gastric cardia tissues, and that esophageal adenocarcinomas (EAC) likely arise from a subset of undifferentiated BE cells expressing both intestinal and stem cell markers [69]. In colon cancer two distinct pre-cancerous states have been identified, with colonic adenomas emerging from the aberrant expansion of normal stem cells, and serrated polyps (precursors for microsatellite-unstable colorectal adenocarcinoma) arising from regions of ‘gastric metaplasia’, comprising cells with absorptive-lineage patterns, gastric gene signatures, and an activated immune environment [70]. Notably, not all metaplastic cells are alike, and different lineages of metaplastic cells may harbor differing levels of cancer risk even in the same patient. Reflecting such lineage heterogeneity, we identified a subgroup of IM cells marked by expression of genes normally expressed in intestinal stem cells (*OLFM4*) (‘intestinal stem-cell dominant’) and another IM subgroup displaying a more differentiated enterocyte phenotype. Single-cell and spatial analysis supports a close relationship between ‘intestinal stem-cell dominant’ IM cells and eventual GC. We propose that similar to BE and EAC, gastric IMs with a higher proportion of intestinal stem-cell dominant IM lineages may be more undifferentiated and harbor a cellular reservoir for the eventual emergence of GC.

One notable finding was the identification of a distinct expression-based molecular subtype of body-resident IMs exhibiting ‘pseudo-antralization’. Pseudo- antralized IMs exhibited both depletions in body/cardia cell types but also increased proportions of antral cell lineages. When contextualized against the existing literature, pseudo-antralized IMs appear to exhibit many previously-described features of SPEM, where aberrant antral type glands form in the stomach body due to parietal cell loss [12] and chief cell transdifferentiation [63]. The molecular distinctiveness of pseudo-antralized IMs was reinforced at the genomic level, as pseudo-antralized IMs exhibited molecular features similar to antral IMs (eg increased clone sizes and mutation rates), but were also distinct from antral IMs exhibiting an elevated *ARID1A* mutation rates and association with incomplete histology. We also found that pseudo-antralized IMs exhibited pronounced inflammatory signatures, potentially implicating chronic inflammation in the pathogenesis of this particular IM subtype. Notably, by analysing IM transcriptomes for microbial sequence reads, we discovered that pseudo-antralized IMs exhibited increased bacterial levels compounded with reduced diversity, a hallmark of microbial dysbiosis linked to multiple gastrointestinal pathologies[71]. Intriguingly, pseudo-antralized IMs were associated with a specific community of microbes normally associated with the healthy oral tract such as *Peptostreptococcus*, *Streptococcus*, and *Prevotella*. A functional role for oral bacteria in the pathogenesis of IM and GC has been recently proposed [72, 73], and lending credence to our results it is worth noting that the oral microbes identified in our study displayed a strong overlap with IM-associated communities defined by more traditional 16S- based sequencing approaches [53]. At the translational level, a role for microbial dysbiosis in IM development may suggest potential interventions for inhibiting the progression of pseudo-antralized IMs through tailored antibiotics or improvements in oral hygiene.

Finally, our findings may have relevance for the management of patients with pre-malignant gastric lesions. Unlike countries such as Japan and South Korea where GC incidence is sufficiently high to warrant unselected population-based screening, mass population screening is not cost-effective in countries where GC incidence is moderate such as Singapore [6]. As an alternative, applying differentiated screening approaches to patients stratified by distinct patterns of GC risk may represent a more sustainable strategy. We have previously reported clinical risk factors such as older age and positive serum pepsinogen indices as strongly associated with early gastric neoplasia [19]. As molecular alterations are also pivotal to GC pathogenesis [74, 75], we evaluated if combining molecular events with clinical models may improve GC risk stratification. Encouragingly, our results revealed that integrating genomic data into clinical risk stratification model improved risk model accuracy, suggesting the potential utility of genomic testing to identify individuals at very high risk of developing GC. *Supplementary Figure 8* proposes a potential clinical pathway for GC precision prevention, where subjects are first risk-stratified by either clinical criteria or inexpensive non-invasive assays (eg blood tests), and those deemed to be high risk are then offered more expensive endoscopic screening and molecular testing. Such a strategy may balance the tension between surveying large patient populations with the resource-intensive investments required for endoscopic procedures and advanced diagnostic testing including genomic sequencing. Ultimately, our results may facilitate the development of a molecularly-guided risk stratification strategy to identify patients at very high risk of GC, and approaches to intercept GC development.

## Contributors

Study concept and design: KKH, KGY, PT. Acquisition of data: All authors. Analysis and interpretation of data: KKH, HM, TU, TS, RHHC, FZ, KGY, PT. Drafting of the manuscript: KKH, RHHC, FZ, KGY, PT.

## Acknowledgements

We would like to thank all our participants and the investigators from NUH, CGH, SGH, TTSH hospitals, investigators from the Singapore Gastric Cancer Consortium (SGCC), the NUHS Tissue Repository, Duke-NUS Genome Biology Facility (DGBF) as well as the staff of all participating endoscopy centres, laboratories and research institutes for their contributions to the study.

