## Supplementary Methods for "Spatiotemporal Genomic Profiling of Intestinal Metaplasia Reveals Clonal Dynamics of Gastric Cancer Progression"

#### Institutional Review Board statements

Approvals were obtained from multiple institutional review boards, including Domain Specific Review Board (DSRB) of the National Healthcare Group (2000/00329), Centralized Institutional Review Board (CIRB) of Singapore Health Services (2018/3222), Institutional Review Board (IRB) of the National University of Singapore (LH-19-070E) and IRB of Seoul National University Hospital (2005-053-1121). All study subjects provided informed consent prior to their participation in the studies.

#### Selection of genomic targets

Candidate genes for GCEP1000 targeted sequencing were selected from a literature review of candidate genes showing (1) significantly mutated or copy number altered genes in gastrointestinal adenocarcinoma, (2) commonly mutated in gastrointestinal adenocarcinoma, and (3) significantly mutated or copy number altered in pre-malignant, inflamed or normal tissues. A total of 277 human genes were selected. We included 6 Hp genes and ~5000 SNPs distributed across the genome to also identify Hp infection and copy number alterations (CNAs). Agilent SureSelect E-array software was used to design unique RNA baits for the gene panel. Biotinylated RNA baits were synthesized by Agilent for use with the SureSelect Target Enrichment system (Agilent, USA).

#### GCEP1000 bulk DNA and RNA extraction and library preparation

Genomic DNA from tissues and blood samples were extracted using the Wizard Genomic DNA Purification Kit (Promega, Madison, Wisconsin, USA) or QIAamp DNA Mini Kit (Qiagen, Hilden, Germany) according to manufacturer protocols. For samples selected for RNA-seq, genomic DNA and total RNA were extracted simultaneously from tissues using the AllPrep DNA/RNA Micro Kit (Qiagen) according to the manufacturer's protocol.

DNA samples were quantified using Qubit brand range assays (Thermo: Q32853) and qualified using Genomic DNA ScreenTapes on a Tapestation (Agilent, 5067-5365). The target enrichment platform was Agilent SureSelect XT HS2 DNA System with Pre-Capture Pooling (Agilent: G9985A, G9985B, G9985C, G9985D) with a customized tier 2 design. Briefly, 100ng of DNA from each sample was enzymatically fragmented (Agilent: 5191-4080) before end-repair, ligation of adaptors and pre-capture amplification performed using unique dual indexing primer pairs. The yield and size distribution of each sample was checked using D1000 ScreenTapes (Agilent:  5067-5582). 16 samples were pooled in equal amount to 1.5 ug per hybridization with the custom panel following manufacturer instructions. The hybridized DNA samples were captured using streptavidin-coated beads before amplification. The yield and size distribution of the captured samples were analyzed on High Sensitivity ScreenTapes (Agilent: 5067-5584). The libraries were then sequenced on the Illumina Novaseq 6000 equipment (PE150bp), according to manufacturer protocols.

Whole genome sequencing libraries were constructed using the New England Biolabs Nextera kit. The genomic DNA was randomly sheared into short fragments, and the obtained fragments were end-repaired, A-tailed, and further ligated with Illumina adapters. The fragments with adapters were PCR amplified, size selected, and purified. The libraries were checked with Qubit and real-time PCR for quantification and on an Agilent bioanalyzer for size distribution detection. Quantified libraries were pooled and sequenced on the Illumina Novaseq 6000 (PE150bp) according to manufacturer's protocols.

10ng of total RNA was used to create RNA-seq libraries using the SMART-Seq Stranded Kit (Takara Bio USA, Mountain View, California, USA) according to the manufacturer protocols. Library fragment size was determined using the High Sensitivity Kit on the Agilent Bioanalyzer (Agilent Technologies). The libraries were sequenced on an Illumina Novaseq 6000 (PE150bp) according to manufacturer protocols.

**GCEP1000 DNA sequencing analysis**

Targeted sequencing reads were aligned to the human reference genome hs37d5 using BWA MEM [1]. Duplicates were removed with Agilent's AGeNT tool using molecular barcode information. Aligned BAM files were further processed according to GATK [2] Best Practices guidelines. Variant calling was performed with Mutect2 [3] using the parameters "--force-active true --pruning-lod-threshold -4 --max-reads-per-alignment-start 0". Candidate variants with less than 5 supporting reads were removed to retain only highly confident predictions. The functional effects of variant were annotated using Funcotator. Genes under positive selection were identified using dNdScv [4], by analysing the ratio of nonsynonymous to synonymous mutations.

CNAs were analyzed using two approaches, the GATK ACNV workflow and ASCAT [5]. Raw copy ratio and allelic copy ratios at targeted regions were collected for both IM and matched blood samples. For GATK, amplified or lost segments were called using CallCopyRatioSegments with default parameters and according to GATK best practices. Allele-specific copy number profiles were separately generated using ASCAT. For ASCAT, Log R ratio (LogR) and B-allele frequency (BAF) plots were manually inspected to select for highly confident CNAs.

For WGS data, sequencing reads were aligned using BWA MEM and processed using GATK, including duplicate removal using MarkDuplicate, local read realignment and base quality score recalibration. Variant calling was performed using standard Mutect2 commands comparing BAM files for IM samples compared to matched blood samples. Mutational signatures were fitted using the signature.tools.lib R package with stomach-specific substitution as reference.

**Bulk RNA-sequencing analysis**

Sequencing reads were aligned to human reference sequence Hg38 using Hisat2 [6], and gene expression was quantified using Stringtie[7]. Differential gene expression and gene set enrichment analysis was performed using DESeq2[8] and fgsea (https://github.com/ctlab/fgsea) respectively.

To quantify bacterial microbiomes, we applied PathSeq[9] to first remove all reads aligned to the human genome, followed by realignment of the remaining reads (with a minimum clipped read-length of 100 bp) to the NCBI database of bacterial reference genomes. Reads that were unambiguously aligned to specific bacterial genera were retained for analysis and the abundance of each bacteria genus normalized to the total number of sequencing reads aligned to the human genome. To estimate bacterial diversity, we used PathSeq normalized (%) scores which are normalized to microbial genome sizes and the overall bacterial scores. All other comparisons used the absolute numbers of unambiguous reads mapping to genus X/reads mapping to human. Oral bacteria was defined by querying their prevalence in healthy oral cavity and stomach using the mBodyMap[10] database.

**Whole exome sequencing**

DNA samples from patients with concurrent adjacent normal, dysplasia and GC were quantified using Qubit brand range assays (Thermo: Q32853) and qualified using Genomic DNA ScreenTapes on a Tapestation (Agilent, 5067-5365). Target enrichment platform used was Agilent SureSelect XT HS2 DNA System with Pre-Capture Pooling (Agilent: G9985A, G9985B) with Agilent SureSelect Human All Exon V6 (5190-8873). Briefly, 100ng of DNA from each sample was enzymatically fragmented (Agilent: 5191-4080) before end-repair, ligation of adaptors and pre-capture amplification with unique dual indexing primer pair. The yield and size distribution of each sample was check on D1000 ScreenTapes (Agilent: 5067-5582). 8 samples were pooled in equal amount to 1.5 ug per hybridization with the SureSelect Human All Exon V6 probes following manufacturer instruction. Hybridization temperature was set at 62.5°C as recommended in the manufacturer instruction. The hybridized DNA samples were captured using streptavidin-coated beads before amplification. The yield and size distribution of the captured samples were analyzed on High Sensitivity SreenTapes (Agilent: 5067-5584). The libraries were also checked with Qubit and real-time PCR for quantification. The quantified libraries were pooled and sequenced on the illumina Novaseq 6000 equipment (PE150bp), according to manufacturer's protocol.

Exome sequencing reads were aligned to the reference human genome hs37d5 using BWA MEM. Preprocessing steps including duplicate marking, local read realignment and base quality score recalibration were performed using Picard and Genome Analysis Toolkit (GATK) to generate analysis-ready BAM files. Mutect2 was used in paired mode to generate a list of somatic SNVs and indels in the 277 genes used in the GCEP1000 panel. GCs were classified as EBV, MSI, CIN or GS using previously proposed classification systems [11]. EBV-positive tumors were identified by evaluating the number of reads mapping to the NC_007605 EBV genome. MSI status was assessed using MSIsensor2, a tool for detecting microsatellite instability from sequencing data. Copy number alterations in the exome data were identified using the GATK ACNV method. Significant somatic copy number alterations in GC samples were identified using GISTIC2. Hierarchical clustering was performed on tumors using copy number profiles from significant copy number altered regions from GISTIC2. Clusters with higher CNV burden were considered as CIN-positive and the remaining samples were considered as GS tumors.

**Single-cell RNA-sequencing**

Patients with IM undergoing endoscopic biopsies at the National University Hospital or Tan Tock Seng Hospital, Singapore were enrolled after written informed consent was obtained. Tissues were collected in MACS tissue storage solution (Miltenyi Biotec) immediately after biopsy and stored on ice. Tissue processing was performed as previously reported [12]. Samples were processed using enzymatic and mechanical dissociation with a human tumor dissociation kit and the Gentle MACS Octodissociator (Miltenyi Biotec) following the manufacturer's "37_h_TDK_2" program. The dissociated cells were passed through a MACS smartstrainer (70 μm) and incubated with RBC lysis buffer for 5 minutes followed by PBS neutralization. All centrifugation steps were carried out at 300× g for 7 minutes. Dissociated cells were washed twice in PBS + 1% bovine serum albumin (BSA). Live-cell counts were obtained by manual cell counting using 1:1 trypan blue dilution. Cells were concentrated to 800–1,200 live cells/μL and processed for single-cell analysis.

Samples from each patient were processed in a single batch for library preparation. The Chromium Single-Cell 3' Library and Gel Bead Kit (10× Genomics) was used according to the manufacturer's protocols. Briefly, gel bead-based emulsions (GEMs) were generated by combining cells, barcoded single-cell 3' Gel Beads and partitioning oil. 10x barcoded full-length cDNAs generated from GEMs were amplified by PCR. Enriched libraries were enzymatically digested, size selected, and adaptor ligated for sequencing. Quantified libraries were sequenced on a Hiseq4000 (Illumina).

Raw fastq sequencing data for each sample was processed using the Cell Ranger V7.0.0 software (https://support.10xgenomics.com/single-cell-gene-expression/software/downloads/latest#cellrangertab) onto the human hg38 reference genome to generate a gene expression count matrix. Subsequently, Seurat V4.1.0 [13] was utilized to perform basic quality control (QC) filtering. Genes shared by fewer than 3 cells and cells with fewer than 500 or more than 7000 genes were filtered out using the Seurat::subset function. Cells with mitochondrial RNA percentage (MT%) higher than median(MT%)+1.5*SD(MT%) were also filtered. DoubletFinder V2.0.3 [14] was employed for each sample to remove potential doublet cells that can often occur in scRNA-seq data and adversely impact the downstream analysis at the single-cell level. Next, processed samples were integrated using the Seurat::merge function, normalized using Seurat::SCTransform, and scaled and analyzed by principal component analysis (PCA). Using the PCA dimension reductional matrix, Harmony V0.1.0 [15] was performed to remove batch effects that may be present in the data. The data was visualized using the Uniform Manifold Approximation and Projection (UMAP) method, and cells were clustered using the Seurat shared nearest neighbour (SNN) algorithm with the Leiden-graph approach.

Ten gastric tumor scRNA-seq data from our previous study [12] were processed separately, following the same workflow as described in the previous section. Copy number variation (CNV) analysis was performed on each sample using CopyKAT V1.0.8 [16] by setting a pool of matched normal cells as reference normal cells. The predicted aneuploid cells were then sub-clustered into sub-groups in a heatmap based on their Euclidian distances on the CNV matrix, and a phylogenetic neighbour joining (NJ) tree was set up using the R package phangorn [17]. The NJ tree was re-rooted using a defined diploid cell. Based on the clustering heatmap and the NJ tree, the predicted aneuploid cells with less CNV burden and closer to the root in the NJ tree were defined as “early-stage tumor cells”.

The raw count data of the IM cells and the early-stage tumor cells were integrated using the Seurat::merge function. The integrated data was then processed following the same workflow as described earlier. Monocle3 V1.2.9 [18] was used to perform trajectory analysis on this data with default parameters. The embeddings of the CDS object were replaced using the Seurat UMAP embeddings for consistency. The root cells of the CDS object were manually selected in both gastric stem cells and intestinal stem cells.

**Digital spatial profiling**

FFPE blocks was cut into five micrometer sections and mounted on BOND Plus slides (Leica Biosystems, Wetzlar, Germany). H&E staining was performed on one slide on which a pathologist demarcated tumour, normal, stroma, lymphoid aggregates and IM regions. The sequential slide was then processed according to GeoMx Human Whole Transcriptome Atlas (NanoString, Seattle, WA, United States). ROI selection was performed based on immune staining using four markers

(DNA, CD45, PanCK and Smooth Muscle Actin). A subset of ROIs were segmented to generate custom AOIs (CD45+ and CD45- regions). 22 to 95 ROIs/AOIs were selected for each slide. Libraries were constructed using Seqcode reagents (NanoString, Seattle, WA, United States) and sequenced on the Illumina platform.

FASTQ files for DSP profiles were processed to generate count matrices as previously described [19]. Briefly, deduplicated sequencing counts were calculated based on UMI and molecular target tag sequences. Single-probe genes were reported as the deduplicated count value. Count data was processed and normalized using the GeoMxTools R package v2.0. AOIs/ROIs with fewer than 1000 raw reads or the percentage of aligned reads <75% or a sequencing saturation <50% were filtered out of the analysis. The limit of quantitation was estimated as 2 geometric standard deviations of the negative control probes above the geometric mean of the negative control probes. AOIs/ROIs that had only a small percentage (< 5%) of panel genes detected above the quantitation limit were removed, and the genes with a low detection rate (< 10%) among the remaining AOIs/ROIs were filtered out. The datasets were normalized using upper quartile (Q3) normalization. To estimate cell abundances in each AOI/ROI, the SpatialDecon algorithm (v.1.4.3) was employed using safeTME as cell-profile matrix.

To distinguish between intestinal stem-cell dominant IM and enterocyte dominant IM, we used Seurat FindMarker to select the top 500 markers (for IM-enterocyte and IM-stem cells each) from our scRNA-seq data. GSEA was performed using these markers for each IM regions, which were normalized to the average gene expression in histologically normal samples to annotate IM regions as IM-enterocyte dominant or IM-stem cell dominant.

**Clinical model**

Logistic regression analysis was used to predict the risk of dysplasia. The clinical risk stratification model was based on four established clinical risk factors - age, pepsinogen (PG) index, OLGIM score, and smoking status. Molecular test results such as mutation count, clone size, and copy number alteration (CNA), were further incorporated into the clinical model to test for its capability to provide additional information on risk prediction beyond present clinical and histological information. Receiver operating characteristic (ROC) curves were used to present the performance of risk factors, with area under the curve (AUC) values as the performance indicator. All statistical analysis was performed using IBM SPSS Statistics 28 (IBM Corp., Armonk, NY, USA). A p-value of less than 0.05 was considered statistically significant.

**Data availability**

Raw sequencing data, including GCEP1000 panel targeted sequencing, bulk RNA sequencing, whole genome sequencing, single-cell RNA sequencing and digital spatial transcriptomic has been deposited at the European Genome-phenome Archive (EGA) under the accession number EGAS00001007067.

### Method references


1 Li H, Durbin R. Fast and accurate short read alignment with Burrows-Wheeler transform. Bioinformatics 2009;**25**:1754-60.

2 McKenna A, Hanna M, Banks E, Sivachenko A, Cibulskis K, Kernytsky A*, et al.* The Genome Analysis Toolkit: a MapReduce framework for analyzing next-generation DNA sequencing data. Genome Res 2010;**20**:1297-303.

3 Cibulskis K, Lawrence MS, Carter SL, Sivachenko A, Jaffe D, Sougnez C*, et al.* Sensitive detection of somatic point mutations in impure and heterogeneous cancer samples. Nat Biotechnol 2013;**31**:213-9.

4 Martincorena I, Raine KM, Gerstung M, Dawson KJ, Haase K, Van Loo P*, et al.* Universal Patterns of Selection in Cancer and Somatic Tissues. Cell 2017;**171**:1029-41 e21.

5 Van Loo P, Nordgard SH, Lingjaerde OC, Russnes HG, Rye IH, Sun W*, et al.* Allele-specific copy number analysis of tumors. Proc Natl Acad Sci U S A 2010;**107**:16910-5.

6 Kim D, Langmead B, Salzberg SL. HISAT: a fast spliced aligner with low memory requirements. Nat Methods 2015;**12**:357-60.

7 Pertea M, Pertea GM, Antonescu CM, Chang TC, Mendell JT, Salzberg SL. StringTie enables improved reconstruction of a transcriptome from RNA-seq reads. Nat Biotechnol 2015;**33**:290-5.

8 Love MI, Huber W, Anders S. Moderated estimation of fold change and dispersion for RNA-seq data with DESeq2. Genome Biol 2014;**15**:550.

9 Walker MA, Pedamallu CS, Ojesina AI, Bullman S, Sharpe T, Whelan CW*, et al.* GATK PathSeq: a customizable computational tool for the discovery and identification of microbial sequences in libraries from eukaryotic hosts. Bioinformatics 2018;**34**:4287-9.

10 Jin H, Hu G, Sun C, Duan Y, Zhang Z, Liu Z*, et al.* mBodyMap: a curated database for microbes across human body and their associations with health and diseases. Nucleic Acids Res 2022;**50**:D808-D16.

11 Cancer Genome Atlas Research N. Comprehensive molecular characterization of gastric adenocarcinoma. Nature 2014;**513**:202-9.

12 Kumar V, Ramnarayanan K, Sundar R, Padmanabhan N, Srivastava S, Koiwa M*, et al.* Single-Cell Atlas of Lineage States, Tumor Microenvironment, and Subtype-Specific Expression Programs in Gastric Cancer. Cancer Discov 2022;**12**:670-91.

13 Hao Y, Hao S, Andersen-Nissen E, Mauck WM, 3rd, Zheng S, Butler A*, et al.* Integrated analysis of multimodal single-cell data. Cell 2021;**184**:3573-87 e29.

14 McGinnis CS, Murrow LM, Gartner ZJ. DoubletFinder: Doublet Detection in Single-Cell RNA Sequencing Data Using Artificial Nearest Neighbors. Cell Syst 2019;**8**:329-37 e4.

15 Korsunsky I, Millard N, Fan J, Slowikowski K, Zhang F, Wei K*, et al.* Fast, sensitive and accurate integration of single-cell data with Harmony. Nat Methods 2019;**16**:1289-96.

16 Gao R, Bai S, Henderson YC, Lin Y, Schalck A, Yan Y*, et al.* Delineating copy number and clonal substructure in human tumors from single-cell transcriptomes. Nat Biotechnol 2021;**39**:599-608.

17 Schliep KP. phangorn: phylogenetic analysis in R. Bioinformatics 2011;**27**:592-3.

18 Cao J, Spielmann M, Qiu X, Huang X, Ibrahim DM, Hill AJ*, et al.* The single-cell transcriptional landscape of mammalian organogenesis. Nature 2019;**566**:496-502.

19 Merritt CR, Ong GT, Church SE, Barker K, Danaher P, Geiss G*, et al.* Multiplex digital spatial profiling of proteins and RNA in fixed tissue. Nat Biotechnol 2020;**38**:586-99.
