## Supplementary Figures for "Spatiotemporal Genomic Profiling of Intestinal Metaplasia Reveals Clonal Dynamics of Gastric Cancer Progression"

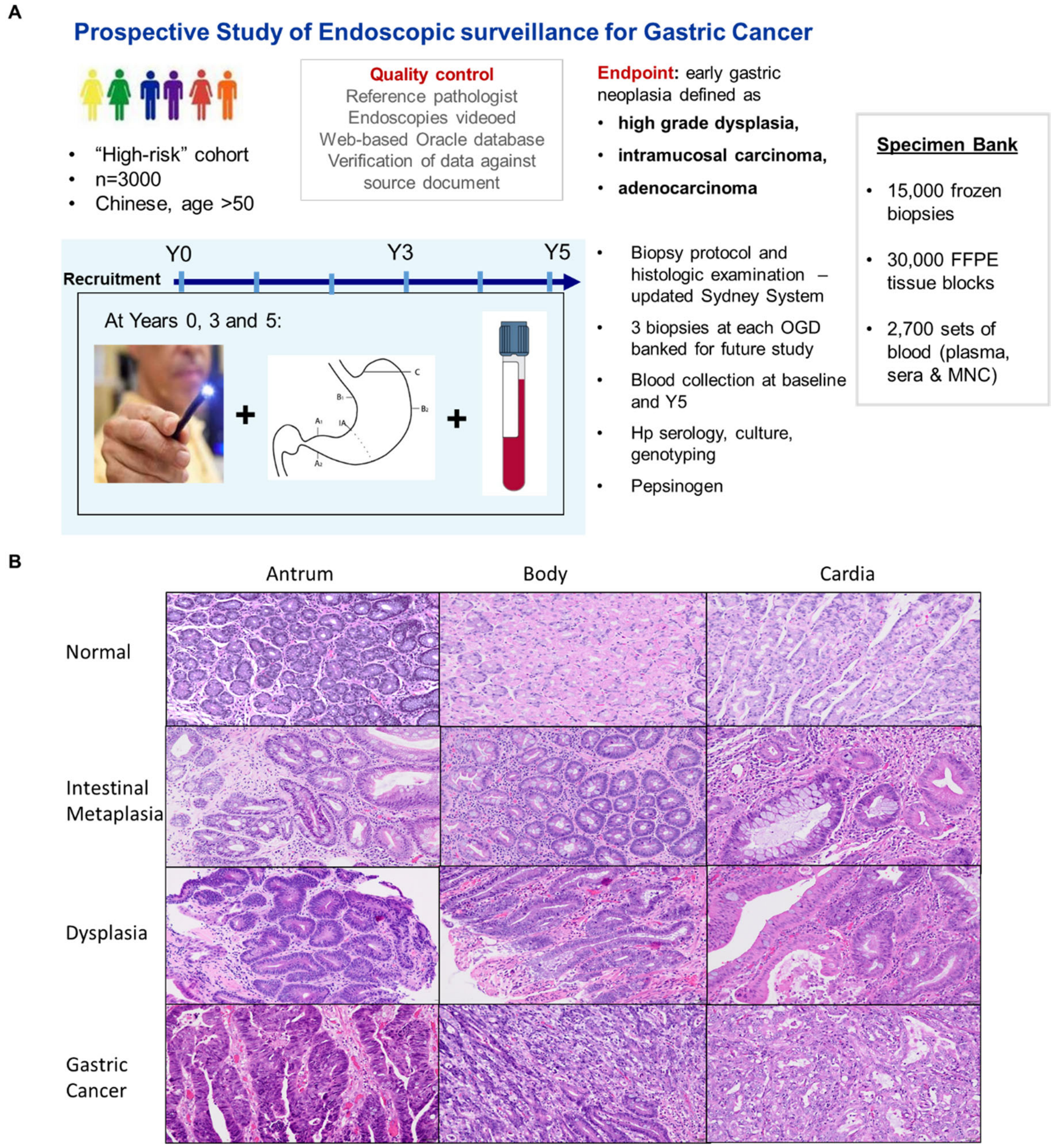


**Figure S1. Overview of Gastric Cancer Epidemiology Program (GCEP).** (A) Quality control metrics (shown) implemented to ensure accurate diagnoses. Samples were collected at years 0, 3, and 5 (multiple samples per patient). Figure is adapted from Huang et al. (2018). (B) Representative H/E images for normal, IM, dysplasia and GC samples from gastric antrum, body and cardia (200X).

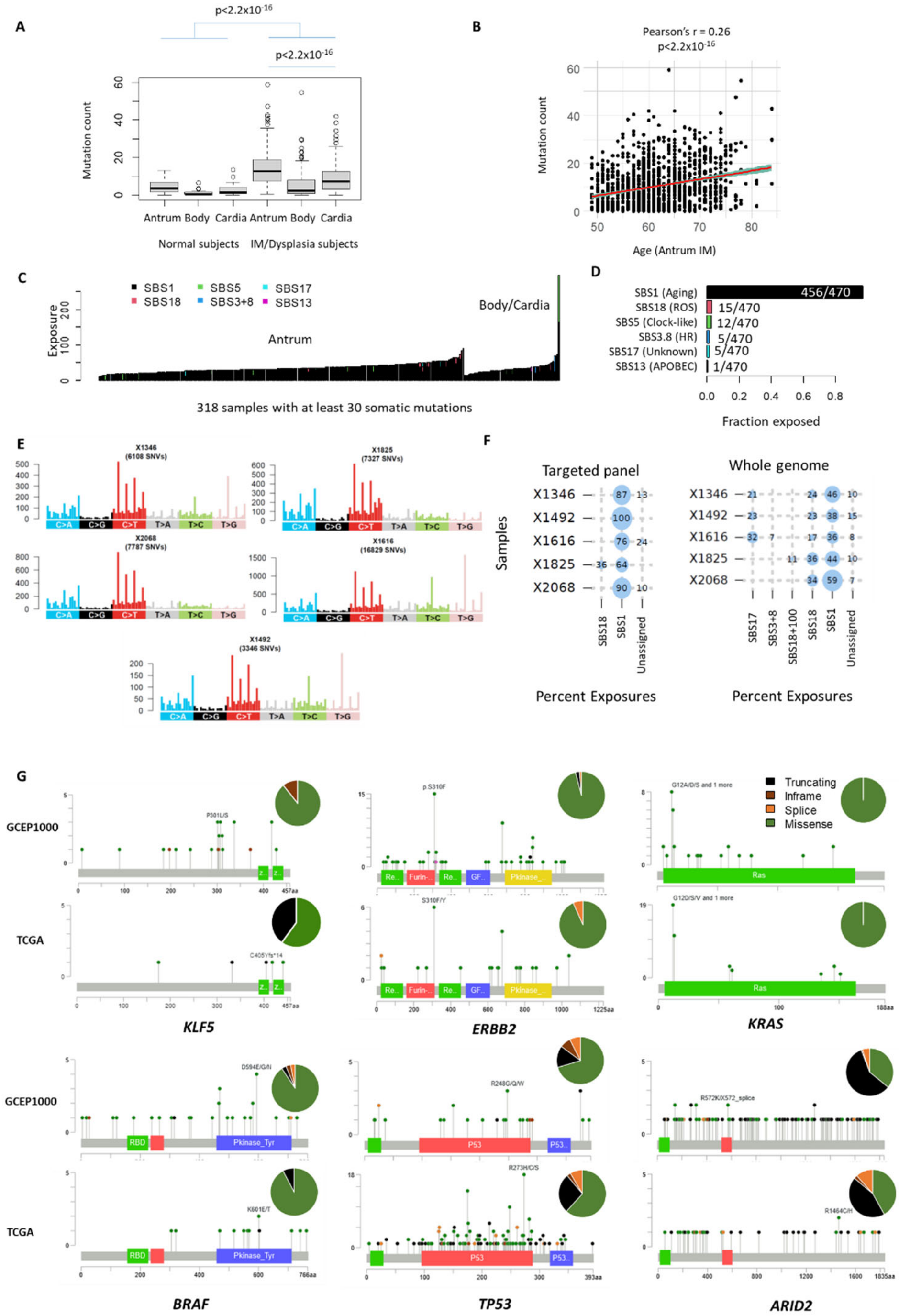


**Figure S2. Genomic landscape of IM and normal samples.** (A) Mutation rate of IM and normal samples at different sites. (B) Correlation between antrum IM mutation count and age. (C) Mutation signatures detected from IM samples. (D) Fraction of IM samples with detectable mutational signatures. (E) Mutation spectrum from 5 antral IM profiled on whole genome sequencing. (F) Proportional exposures to mutation signatures in 5 antral IM from GCEP1000 targeted panel and whole genome sequencing. (G) Lollipop plots highlighting genomic alterations in selected genes in pre-malignant IM (GCEP) and gastric cancer (TCGA)

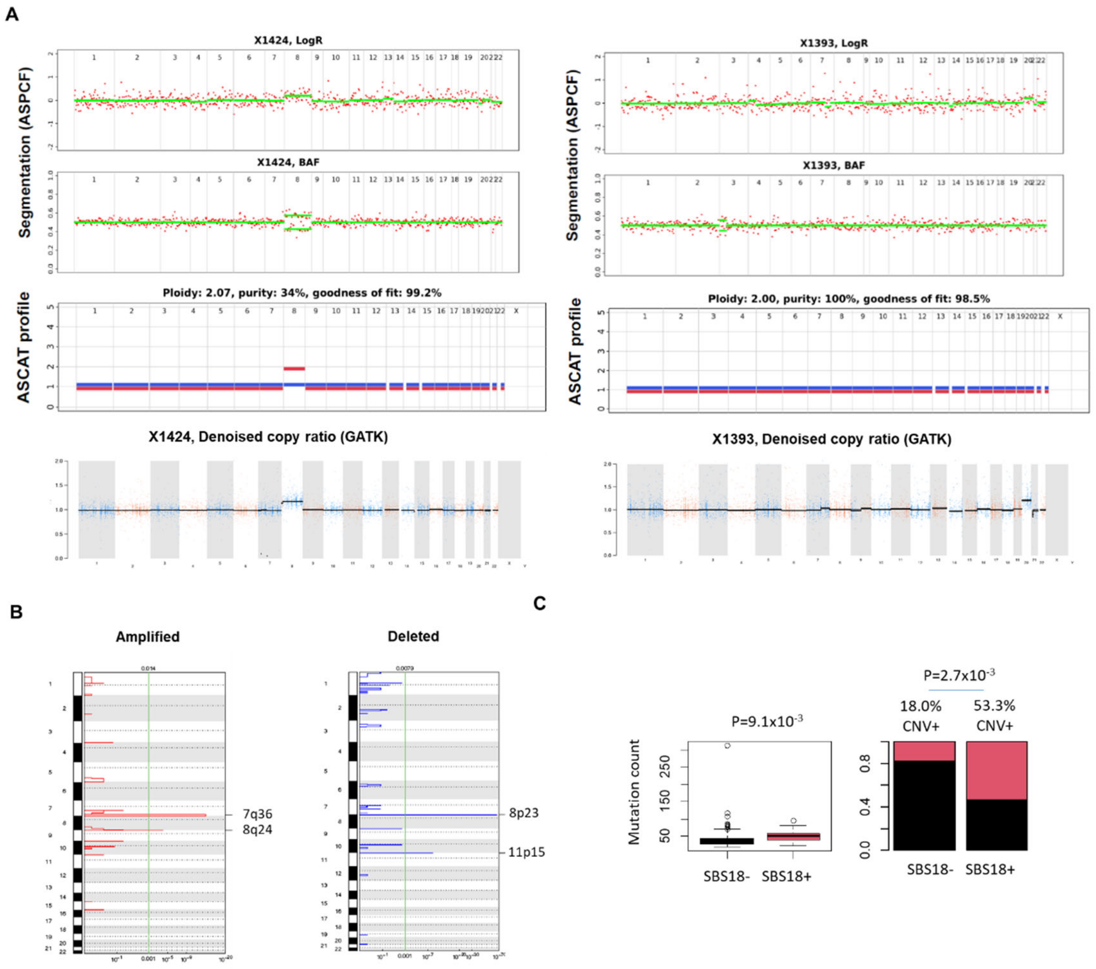


**Figure S3. Copy number landscape of IM.** (A) Example of a CNA identified using both ASCAT (logR and BAF) and GATK (left); and another CNA detected using GATK only (right). For X1393, ASCAT plots show increased logR depths (chromosome 20) but the BAF remains flat, which may represent CNAs at very low purity or sequencing noise. (B) GISTIC amplification and deletion regions of CNA regions identified by GATK (C) Mutation count and CNA presence in samples with and without detectable SBS18 signatures from targeted panel.

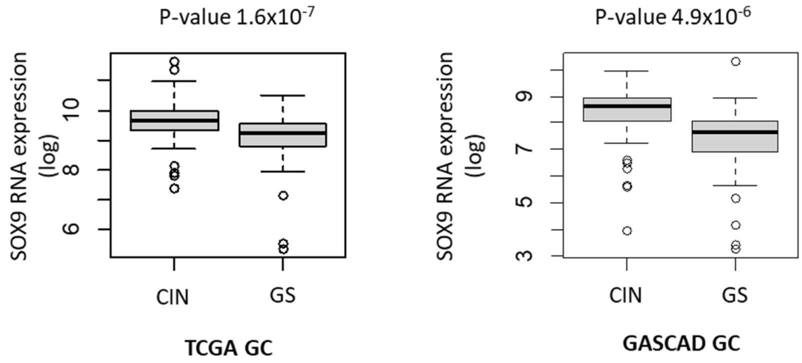


**Figure S4. SOX9 expression in CIN vs GS TCGA (left) and GASCAD (right) GC samples.**

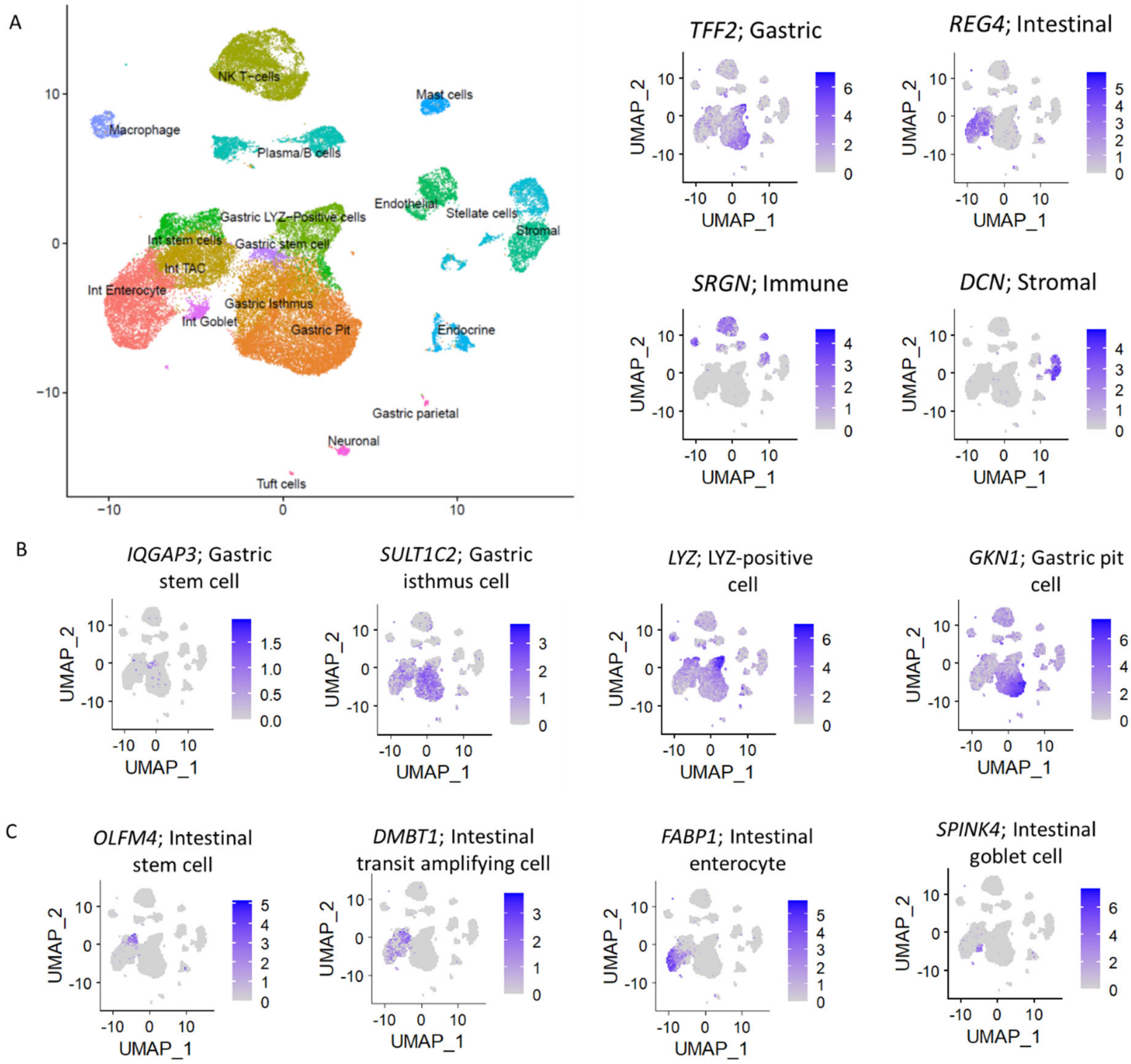


**Figure S5. Annotation of single-cell RNAseq clusters.** (A) Marker genes for gastric (*TFF2*), intestinal (*REG4*), immune (*SRGN*) and stromal (*DCN*) lineage clusters. (B) Marker genes for gastric lineage cell clusters. (C) Marker genes for intestinal lineage cell clusters.

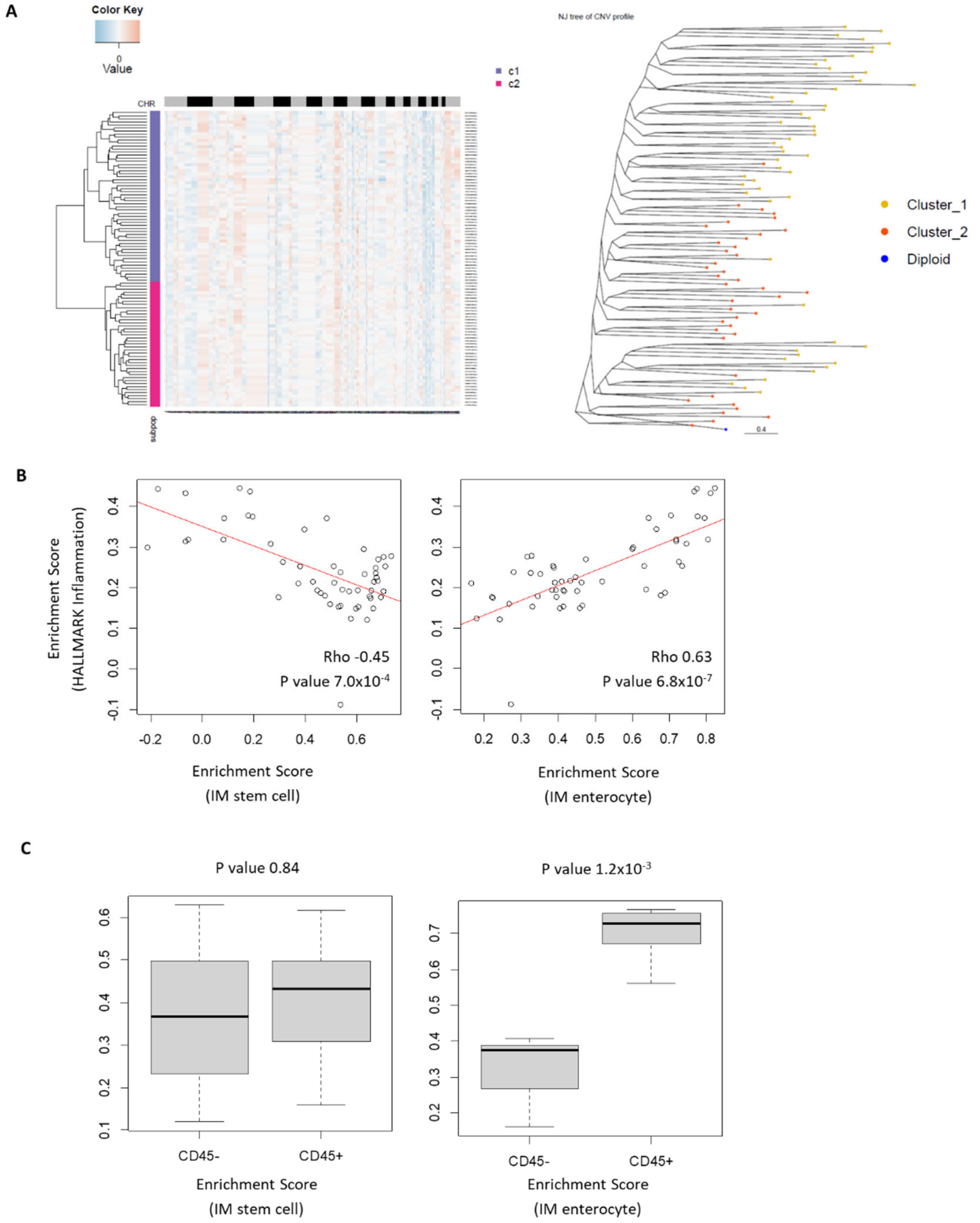


**Figure S6. Comparison between IM and GC.** (A) Copy number inference in a GC patient epithelial single cells (left). Neighbour joining clustering of single cells using a diploid cell as root. Cell cluster with smaller copy number burden and cluster closer to diploid cell is considered as early stage GC. (B) Spearman correlation between IM stem cell (left) and enterocyte (right) enrichment score in DSP ROIs with the HALLMARK inflammation enrichment score. (C) Enrichment score of IM stem cell (left) and enterocyte (right) in CD45- and CD45+ AOIs.

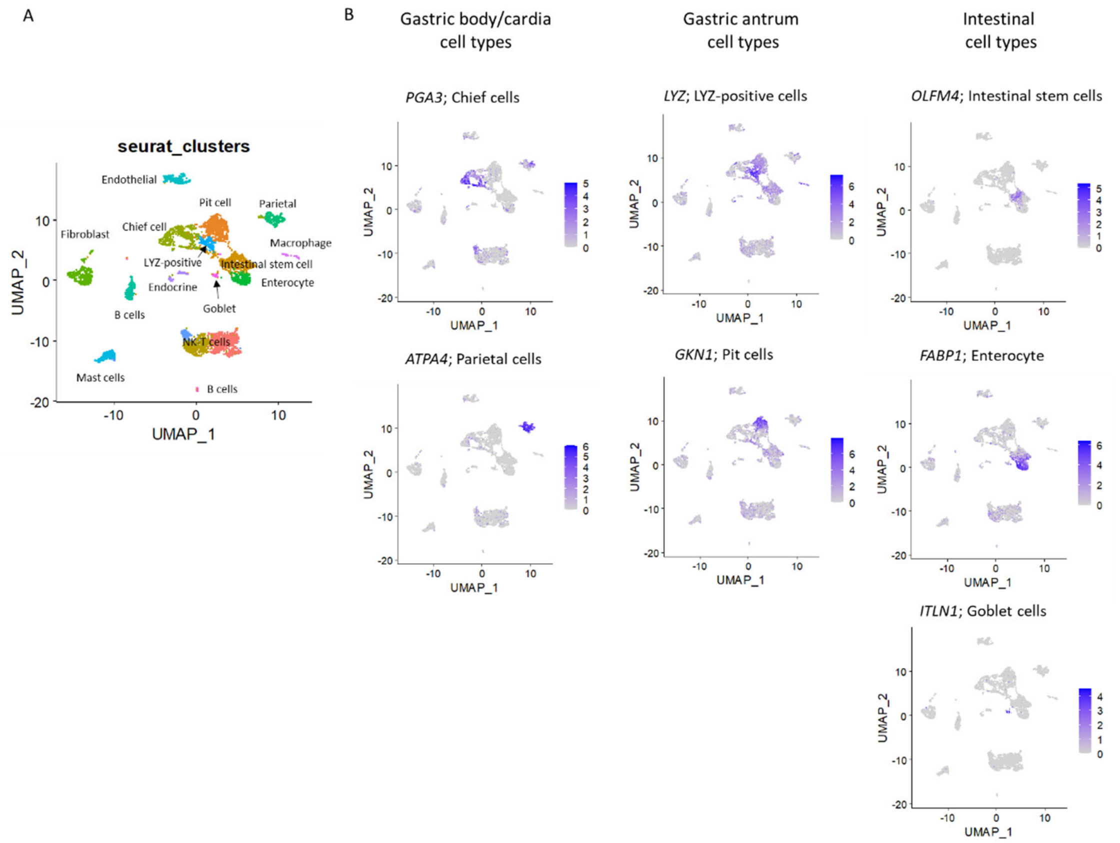


**Figure S7. Single cell RNAseq profile in gastric body biopsies.** (A) Cell clusters identified from 4 gastric body samples (3 IMs and 1 normal). (B) Marker genes used to annotate cell types in antrum, body/cardia and intestine.

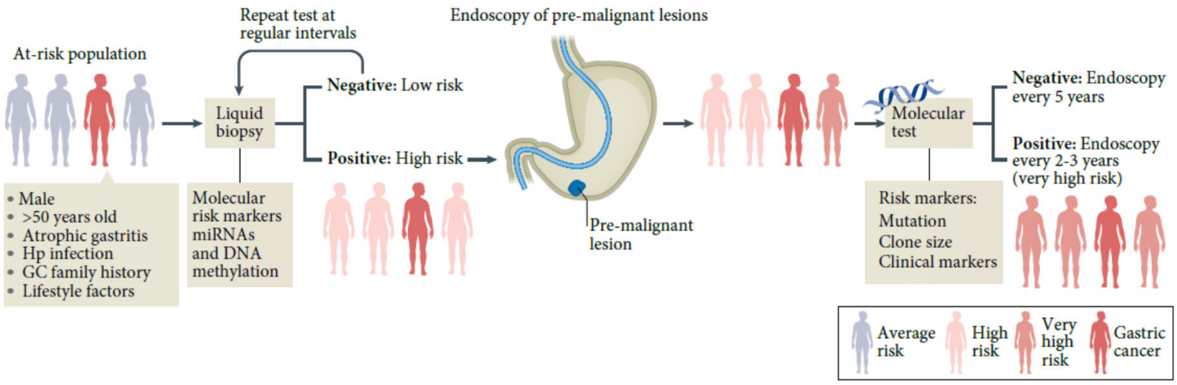


**Figure S8. Precision prevention strategies for GC.** In regions of low and intermediate gastric cancer (GC) risk, regular endoscopic screening is cost prohibitive. Surveillance of patients with pre-malignant conditions, such as intestinal metaplasia, using molecular tests detecting mutation load and genetic clones may be useful in stratifying 'very-high-risk' individuals for intensive and repeated endoscopic follow-up. Figure is adapted from Yeoh and Tan (2022).
